# Deconvolution of bulk blood eQTL effects into immune cell subpopulations

**DOI:** 10.1101/548669

**Authors:** R. Aguirre-Gamboa, N. de Klein, J. di Tommaso, A. Claringbould, U. Võsa, M. Zorro, X. Chu, O.B. Bakker, Z. Borek, I. Ricaño-Ponce, P. Deelen, C.J. Xu, M. Swertz, I. Jonkers, S. Withoff, I. Joosten, S. Sanna, V. Kumar, H.J.P.M. Koenen, L.A.B. Joosten, M.G. Netea, C. Wijmenga, BIOS Consortium, L. Franke, Y. Li

## Abstract

Expression quantitative trait loci (eQTL) studies are used to interpret the function of disease-associated genetic risk factors. To date, most eQTL analyses have been conducted in bulk tissues, such as whole blood and tissue biopsies, which are likely to mask the cell type context of the eQTL regulatory effects. Although this context can be investigated by generating transcriptional profiles from purified cell subpopulations, the current methods are labor-intensive and expensive. Here we introduce a new method, *Decon2*, a statistical framework for estimating cell proportions using expression profiles from bulk blood samples (Decon-cell) and consecutive deconvolution of cell type eQTLs (Decon-eQTL). The estimated cell proportions from Decon-cell agree with experimental measurements across cohorts (R ≥ 0.77). Using Decon-cell we can predict the proportions of 34 circulating cell types for 3,194 samples from a population-based cohort. Next we identified 16,362 whole blood eQTLs and assign them to a cell type with Decon-eQTL using the predicted cell proportions from Decon-cell. Deconvoluted eQTLs show excellent allelic directional concordance with those of eQTL(≥ 96%) and chromatin mark QTL (≥87%) studies that used either purified cell subpopulations or single-cell RNA-seq. Our new method provides a way to assign cell type effects to eQTLs from bulk blood, which is useful in pinpointing the most relevant cell type for a certain complex disease. Decon2 is available as an R package and Java application (https://github.com/molgenis/systemsgenetics/tree/master/Decon2), and as a web tool (www.molgenis.org/deconvolution).

## Introduction

For many of the genetic risk factors that have been associated to immune diseases by genome-wide association studies (GWAS), the molecular mechanism leading to disease remains unknown^1^. Most of these genetic risk variants are located in the non-coding regions of the genome, implying that they play a role in gene regulation^2,3^. Expression quantitative trait locus (eQTL) analysis provides a way to characterize the regulatory effect of these risk factors in humans, and many eQTL studies have now been carried out using bulk tissues, for example, whole blood^4,5^. However, bulk tissues comprise many different cell types, and gene regulation is known to vary across cell types^6–8^. In recent years, efforts to describe eQTL effects in purified cell subpopulations have been carried out in specific cell types^9^. Unfortunately, the length and cost of the study protocols have limited these studies to small sample sizes and only a few cell types. Nevertheless, being able to pinpoint the particular cell type (CT) in which a risk factor exerts an eQTL effect could help us to understand its role in disease.

Statistical approaches to detect CT effects using tissue expression profiles have mainly been developed to evaluate gene by environment interaction (GxE) terms, for example, being able to detect CT eQTLs for myeloid and lymphoid lineages using only whole blood gene expression and by evaluating the interaction between genotype and cell proportions for neutrophils and lymphocytes in whole blood^10^. A second study linked eQTL genes to proxy genes through correlation; these proxy genes were then associated to intrinsic or extrinsic factors, such as cell proportions or inflammation markers^11^. However, these efforts focused on exploiting only one GxE term, or on indirectly linking the CT proportions to given eQTL instead of directly ascertaining the interaction between all the main cell proportions comprising the bulk tissue and genotype. Unfortunately, quantifying cell proportions, in particular for rare subpopulations (total abundance of ≤ 3% in circulating white blood cells), is expensive and time-consuming. Hence, quantifying immune cell proportions in large functional genomics cohorts is not common practice.

Here we present and validate Decon2, a computational and statistical framework that can: (1) predict the proportions of known circulating immune cell subpopulations (Decon-cell), and (2) use these predicted proportions along with whole blood gene expression and genotype information to assign bulk eQTL effects into CT eQTLs (Decon-eQTL). Our two-step framework provides an improvement over previously published methods. As unlike earlier methods^12^, Decon-cell does not rely on any prior information of transcriptome profiles from purified cell subpopulations, as it only requires the proportions of the cells comprising the bulk tissue, in this case whole blood, and identifies signature genes which correlate with cell proportions in a bulk tissue. Secondly, Decon-eQTL is the first approach in which all major cell proportions (the major cell types for which the sum of proportions per sample to approximately 100%) of bulk blood tissue are incorporated into an eQTL model simultaneously. This can then be systematically tested for any significant interaction between each CT and genotype, while at the same time the effect of the other CTs are modelled.

We generated the Decon-cell predictive models using data from the 500FG cohort^13^, where quantification of immune cell types was carried out using FACS^14^ and RNA-Seq based bulk whole blood transcriptome profiling were available for 89 samples^15^. By using a cross-validation approach we were able to accurately predict 34 out of 73 cell subtypes using solely whole blood gene expression. For validation, we applied Decon-cell to three independent cohorts (Lifelines Deep^16^, n = 627, Leiden Longevity cohort^17^, n= 660 and the Rotterdam Study^18^, n= 773) with both blood RNA-seq and measured cell proportion data available (neutrophils, lymphocytes and CD14+ monocytes and granulocytes). Additionally, we benchmarked Decon-cell prediction performance against two other existing methods that quantify immune cell composition using gene expression profiles from whole blood. After showing that we can accurately predict circulating immune cell proportions, we applied Decon-cell to estimate cell proportions in 3,194 individuals from the BIOS cohort^16,19–21^, in which both whole blood RNA-seq and genotypes were available. The BIOS cohort is a valuable resource for functional genomics studies where extensive characterization of the genetic component on gene expression ^11^ and epigenetics^22^ have been performed. We integrated whole blood expression, genotype information and predicted cell proportion with Decon-eQTL, to deconvolute 16,362 significant whole blood cis-eQTLs top effects into CT eQTLs. These deconvoluted eQTL results were comprehensively validated using transcriptome profiles from purified cell subpopulations^23^, eQTLs and chromatin mark QTLs from purified cell types^9^, and eQTLs from single-cell experiments^24^. We also systematically compared the performance of Decon-eQTL against previously published methods^10–11^ that detect cell type eQTL effects using whole blood expression profiles.

## Results

### Decon-cell accurately predicts the proportions of known immune cell types

In order to assign the CT in which an overall eQTL effect from a bulk tissue sample (e.g. whole blood), we need three types of information: genotype data, tissue expression data, and cell type proportions (**Fig. 1**). Here, we propose a computational method for predicting the cell proportions of known immune cell types using gene signatures in whole blood expression data by employing a machine-learning approach. Decon-cell employs the regularized regression method elastic net^25^ to define sets of signature genes for each cell type. In other words, these signatures were selected as having the best prediction power for individual cell proportions.

**Figure 1.**
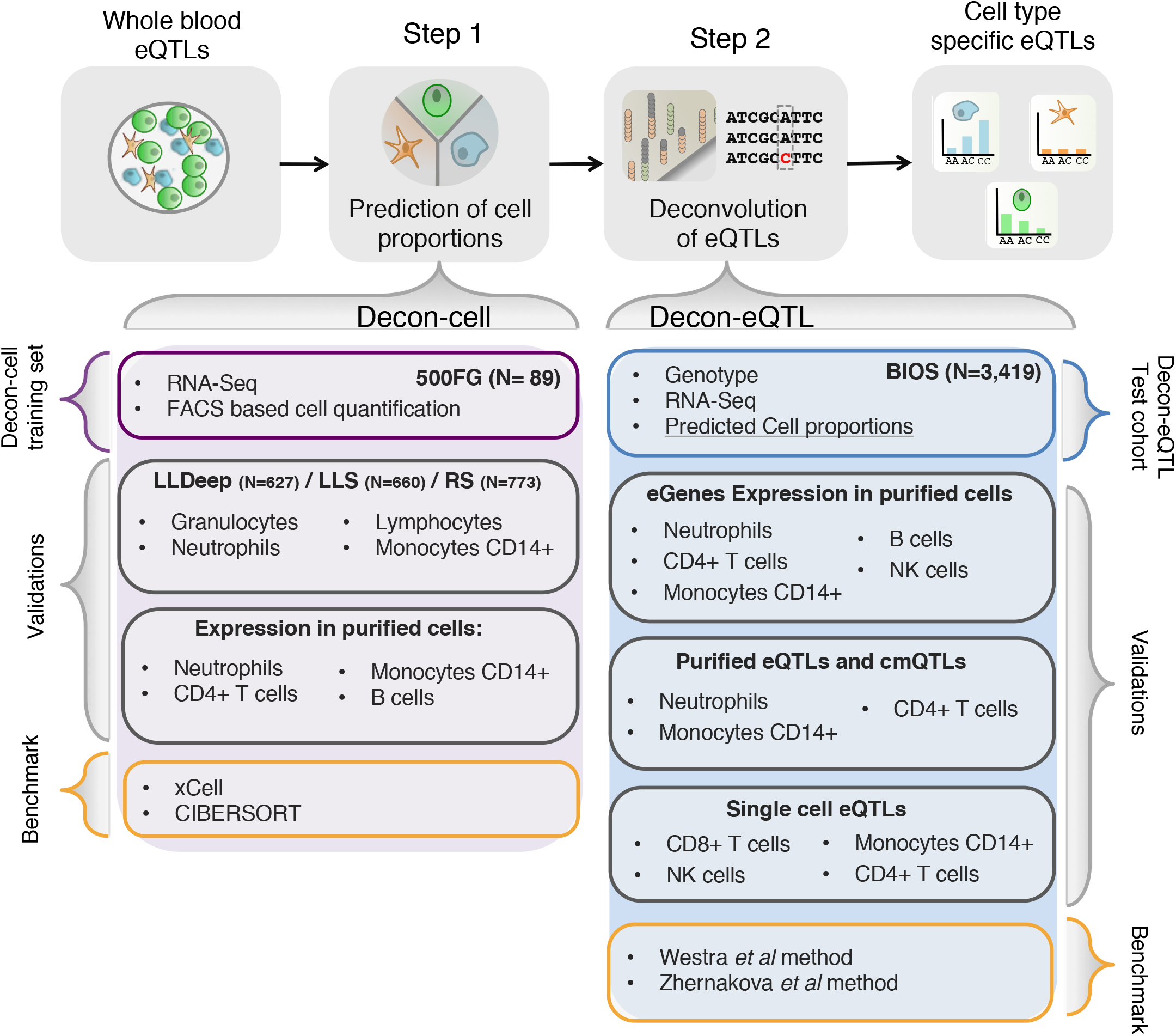
Workflow of application of Decon2 to predict cell counts followed by deconvolution of whole blood eQTLs. With whole blood expression and FACS data of 500FG samples, Decon-cell predicts cell proportions with selected marker genes of circulating immune cell subpopulations. Validations of Decon-cell were carried out on three independent cohorts where measurements of neutrophils/granulocytes, lymphocytes and monocytes CD14+ were available, alongside to expression profiles of whole blood. Benchmarking of Decon-cell was performed against CIBERSORT^26^ and xCell^12^. Decon-cell was applied to an independent cohort (BIOS) to predict cell counts using whole blood RNA-seq. Decon-eQTL subsequently integrates genotype and tissue expression data together with predicted cell proportions for samples in BIOS to detect cell type eQTLs. We validated Decon-eQTL using multiple independent sources, including expression profiles of purified cell subpopulations, eQTLs and chromatin mark QTLs (cmQTLs) from purified neutrophils, monocytes CD14+ and CD4+ T cells^9^, and single cell eQTLs results^24^. Benchmarking of Decon-eQTL was carried out for comparison with previously reported methods which detected cell type eQTL effects using whole blood expression data, i.e. Westra method ^10^ and Zhernakova, *et al* method^11^).

There are 89 samples in 500FG cohort with both whole blood RNA-seq and quantification of 73 immune cell subpopulations by FACS available. This data was used to build the prediction models for estimating cell subpopulations by Decon-cell. First we determined which of the 73 cell subpopulations could be reliably predicted by Decon-cell. A within-cohort cross-validation strategy was employed by randomly dividing 89 samples (**Fig. 1**) into training and test sets (70% and 30% of the samples, respectively). After generating a model using each training set, we applied the prediction models of each cell type to the samples in the test sets. We compared the predicted and measured cell proportion for each cell type using Spearman correlation coefficients to evaluate the prediction performance. We repeated this process 100 times and then used the mean correlation coefficient in all 100 iterations to evaluate the prediction performance. We were able to predict 34 out of 73 cell subpopulations using whole blood gene expression data at a threshold of mean absolute R ≥ 0.5 across all 100 iterations (**Fig. 2A, Supplementary Fig.1**, Supplementary Table 1). The number of signature genes selected in the models for predicting cell proportions varied across the cell types, ranging from 2 to 217 signature genes (Supplementary Fig. 2A, Supplementary Table 1); and it was independent of the average abundance of these cell types in whole blood (R = 0.02, Spearman correlation coefficient, Supplementary Fig.2A). In particular, cell types that are abundant in whole blood (granulocytes-neutrophils, CD4+ T-cells, CD14+ monocytes) were predicted with high confidence (correlation between predicted and measured values, R ≥ 0.73). Remarkably, we were also able to predict a number of less abundant cell subpopulations, including NK cells, CD8+ T-cells, non-NK T-cells (CD3-CD56-) including CD4+ central memory and CD4+ effector memory T-cells, and regulatory T-cells (Supplementary Fig. 2A) as determined by FACS. Cell types with a low prediction performance (R < 0.5) are those that have few signature genes whose expression levels correlate sufficiently (i.e. absolute R < 0.3) with the actual cell proportions in whole blood (Supplementary Fig. 2B-C). For each of the 34 predictable cell types, we used Decon-cell to build models for predicting their cell counts using all 89 samples from the 500FG cohort. These models were applied to 3,194 samples in an independent cohort, to predict cell proportions of circulating immune cell types for the subsequent deconvolution of eQTL effect.

**Figure 2.**
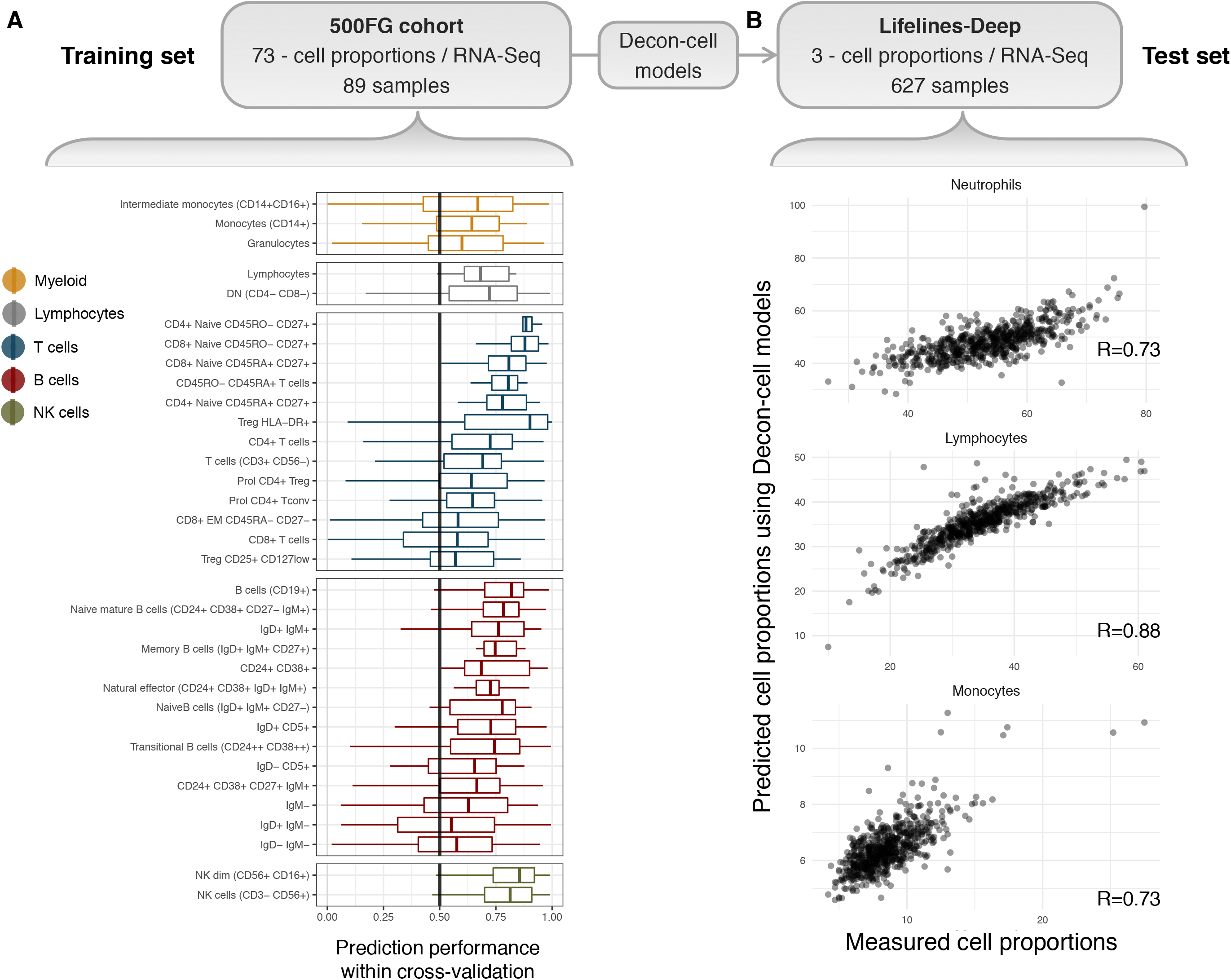
Prediction of cell proportions using whole blood transcriptome by Decon-cell. (A) Distribution of prediction performance (Spearman correlation coefficient) of the 34 predictable cell types in 100 iterations of prediction within the 500FG cohort. (B) Cross-cohort validation in an independent Lifelines-Deep cohort (n=627): the measured and predicted cell proportions for neutrophils (given by granulocytes in 500FG), lymphocytes and monocytes are compared.

In addition to within-cohort validation, we tested our cell proportion models using three independent cohorts (LLDeep, n = 627, LLS, n= 660, RS, n =773), for which cell type abundances were quantified using a Coulter counter for neutrophils (granulocytes for RS), lymphocytes, and CD14+ monocytes (**Fig. 2B, Supplementary Fig. 3A-B**). In LLDeep we were able to accurately predict these three cell types with Spearman correlation coefficients of R = 0.73, R = 0.89, and R = 0.73, respectively. For LLS and RS the prediction performance was also accurate for neutrophils and lymphocytes, but less accurate for monocytes (R= 0.76 for neutrophils, R= 0.50 for CD14+ monocytes and R= 0.84 for lymphocyte proportions in LLS, R= 0.74 for granulocytes, R= 0.28 for CD14+ monocytes and R= 0.83 for lymphocytes in RS).

Next, in order to benchmark Decon-cell we have compared its prediction performance against two other existing tools that quantify the abundance of known immune cell types using bulk whole blood expression profiles: CIBERSORT^26^ and xCell^12^. We obtained the predicted proportions by CIBERSORT and enrichment scores of circulating immune cells by xCell for the samples in three different cohorts: LLDeep, LLS and RS (Supplementary Fig. 4A-B). For each cell type, Decon-cell outperforms CIBERSORT and xCell (Supplementary Fig. 3B). The scatterplots of predicted vs measured values (Supplementary Fig. 3 A, and Supplementary Fig. 4 A-B) further demonstrate that the better performance of Decon-cell is not due to cell proportion outliers.

Finally, we evaluated whether the signature genes showed CT expression in their relevant purified cell types, using the BLUEPRINT^23^ RNA-seq data from the purified cell subpopulations. We focused on cell types with more than three samples measured, these included neutrophils, CD14+ monocytes, CD4+ T-cells and B-cells. The signature genes showed overall higher expression in their relevant cell subpopulations compared to other cell subpopulations. Interestingly, the signature genes were also able to cluster the samples of the relevant CT using unsupervised hierarchical clustering (Supplementary Fig. 5A-D). Together, our results demonstrated that the gene signatures identified by Decon-cell were predictive for the proportions of circulating immune cell subpopulations using only whole blood gene expression data.

To facilitate the cell proportion prediction of new samples using whole blood RNA-seq, we have made the Decon-cell prediction models and gene signatures available in an R package (Decon-cell) and as a web tool (www.molgenis.org/deconvolution). These two implementations allow the user to pre-process their RNA-seq expression counts and estimate cell proportions using the pre-established models for 34 cell types in whole blood. Decon-cell R package also allows the user to input bulk expression profiles and cell proportions to generate predictive models for new tissues.

### Decon-eQTL assigns bulk eQTLs to cell types eQTL effects

As we know, eQTL analysis using whole blood bulk expression data fails to distinguish between a general eQTL that is present in all cells and an effect that is mainly found in a particular cell subpopulation, or subset of one, significantly more than the others present in the tissue. We therefore propose a new approach to assign the overall bulk eQTL into CT effects, called Decon-eQTL (see **Online Methods**). By using the cell proportions in whole blood, is possible to formally test if the genetic effect is dependent on cell proportions. More explicitly, we include both the genotype and all CT proportions of interest in a linear model and systematically test if there is a significant interaction effect between the genotype and each of the predicted cell proportions in the variation of gene expression in whole blood. At the same time we control the effect of the remaining CTs. In this way, whole blood expression data, alongside genotypes and (predicted) cell proportions can be integrated to assign a CT effect from a bulk eQTL(Fig. 1).

We applied Decon-eQTL to 3,198 samples (BIOS cohort) with transcriptome levels (RNA-seq), genotype information and cell proportions predicted by Decon-cell. Whole blood *cis-e*QTL mapping yielded 16,362 whole blood eQTLs (false discovery rate (FDR) ≤ 0.05). For each of these whole blood *cis-e*QTLs, we applied Decon-eQTL with a focus on 6 major cell subpopulations: granulocytes, CD14+ monocytes, CD4+ T-cells, CD8+ T-cells, B-cells and NK cells. These cell types were selected as the sum of their relative percentages was close to 100% and none of these cell type pairs had an absolute correlation coefficient R ≥ 0.75. Decon-eQTL computationally assigned 4,139 CT eQTLs from these subpopulations, reflecting 3,812 genes and 3,650 SNPs. We observed that 25% of the whole blood eQTLs have a significant (FDR ≤ 0.05) CT eQTL effect given Decon-eQTL. The majority (31%) of the total CT-eQTL effects detected were found to be associated to granulocyte, possibly because granulocytes comprise ∼70% of circulating white blood cells (**Fig. 3A**). We also observed that the majority (74%) of CT eQTLs detected by our method were assigned to a single cell type. It should be noted that these eQTL are likely not exclusively present for this particular cell type in biology, but that the statistical power was sufficient to detect CT eQTL in this particular cell type (Supplementary Fig. 6). We found sharing of eQTLs for between cell types only in few cases. An example of such shared eQTLs is on *NOD2 gene*, where Decon-eQTL was able to detect a strong granulocyte-eQTL effect alongside a smaller, opposite effect in CD14+ monocytes. This opposite effect has also been previously described in eQTL studies on purified CD14+ monocytes and neutrophils^8^. These results demonstrate that cell type effects should be taken into account when interpreting eQTLs derived from bulk tissues.

**Figure 3.**
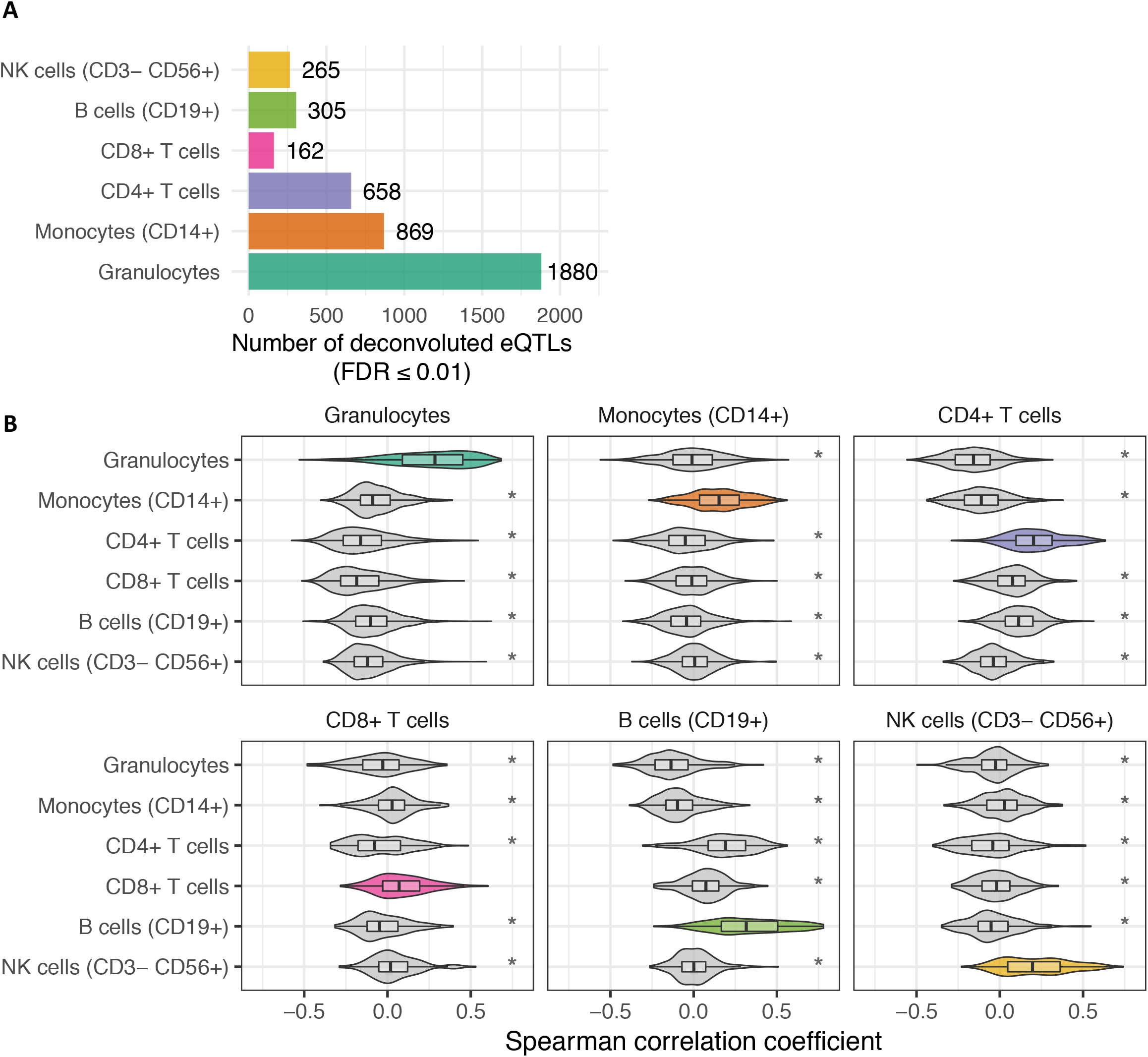
Deconvolution of whole blood eQTLs into cell-type eQTLs. By integrating proportions of cell subpopulations (predicted by Decon-cell), gene expression and genotype information, Decon-eQTL detect cell-type eQTLs. (A) The number of deconvoluted eQTLs in each cell type by using whole blood RNA-seq data of 3,189 samples in BIOS cohort. (B) Distribution of Spearman correlation coefficients between expression levels of deconvoluted eQTL gene and cell counts for each cell subpopulation. The deconvoluted eQTL genes show positive and statistically higher correlation (Spearman) with its relevant cell type proportions than compared to the rest (T test p value < 0.05) in an independent cohort (500FG).

### Decon-eQTL prioritizes genes to relevant cell types

To further validate our deconvoluted CT eQTLs, we systematically tested if the expression levels of the CT eQTL genes detected in the BIOS cohort were correlated with their relevant cell proportions. We calculated the Spearman correlation coefficients between the expression of the identified CT eQTL genes and the measured cell proportions using the 500FG cohort (n = 89). Next, we compared the correlation coefficients obtained with those between expression and the remaining cell proportions. For each of the six evaluated cell subpopulations in Decon-eQTL, CT eQTL genes had a significantly higher correlation with their relevant cell subpopulation than the other cell types (T test, p-value < 0.05) (Fig. 3B). As such, this result validates the association of the CT-eQTL genes and the cell proportions in an independent cohort.

Next, we evaluated whether the significant CT eQTL genes were over-expressed in their relevant cell subpopulation compared to eQTL genes that were found to be non-significant for the same cell type. For this purpose, we made use of the purified neutrophil, CD14+ monocyte, CD4+ T-cell and B-cell RNA-seq data from the BLUEPRINT dataset. We only include these cell types as they were the only ones with more than 3 samples measured. For each of the four cell types, we observed that the expression of CT eQTL genes detected by Decon-eQTL was significantly higher (T-test, p-value ≤ 0.05) compared to the expression of non-significant Decon-eQTL genes (**Fig. 4A**). We also observed that the deconvoluted eQTL genes from granulocytes showed a relatively wider range of variation than the CT-eQTL genes from the other three subpopulations. We hypothesized that this could be explained by the fact that granulocytes comprise ∼70% of the cell composition in whole blood, thus giving us the power to detect eQTL for lowly-expressed genes in granulocytes. This was partly supported by the observation that the variation of expression in whole blood for granulocyte eQTL genes was significantly greater than those eQTL genes deconvoluted to the other five cell subpopulations (F test, p-value ≤ 0.05, Supplementary Fig. 7).

**Figure 4.**
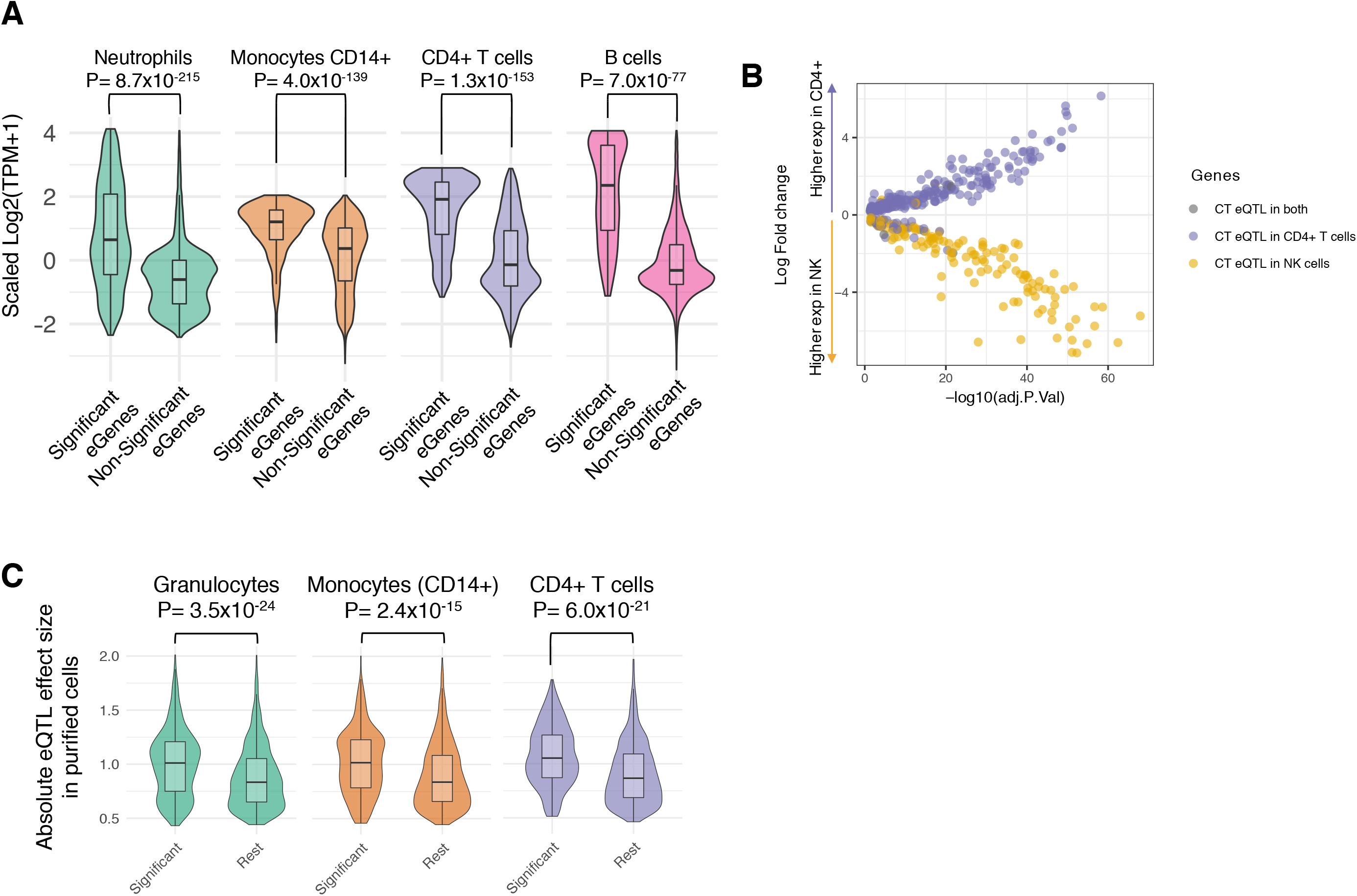
Validation of deconvoluted cell-type eQTLs. (A) Expression of eQTL genes in purified cell subpopulations from BLUEPRINT^23^ is significantly higher in its relevant cell subpopulation compared to other available cell subtypes (green for granulocyte eQTL genes showing expression for purified neutrophils; orange for monocytes; purple for CD4+ T cells; pink for B cells). (B) Differential expressed genes (Adjusted p-value ≤ 0.5) between CD4+ T cells and NK cells are significantly enriched for CT eQTLs effects on CD4+ T cells (dots in purple, Fisher exact P = 1.8×10^17^) and NK Cells (dots in yellow, Fisher exact P = 2.3×10^18^) respectively. (C) Deconvoluted eQTLs (FDR ≤ 0.05) show significantly larger effect sizes in the purified cell eQTLs data ^9^ compared to the rest of the whole blood eQTLs for which we do not detect cell type effect, as shown for deconvoluted granulocyte eQTLs in neutrophil derived eQTLs (green); monocytes (orange); CD4+ T cells (purple).

Furthermore, by using a publicly available transcriptome profiles (GSE78840^27^) of purified NK cells and CD4+ T cells, we assessed if the differentially expressed genes across the two cell types were enriched for eGenes of deconvoluted CT eQTLs. We observed that the CD4+ differentially expressed genes (Adjusted P-value ≤ 0.05) were significantly enriched for CD4+ T cell eQTLs (Fisher exact P = 1.8×10^−17^), whereas NK cell differential genes (Adjusted P-value ≤ 0.05) were significantly enriched for NK cell eQTLs (Fisher exact P = 2.3×10^−18^) as shown in **Fig.4B**.

In summary, we were able to show that the eQTL genes detected by Decon-eQTL have a relevant cell-type effect given we have been able to show that transcriptionally active in their relevant cell type

### CT eQTLs identified by Decon-eQTL in whole blood are replicated in purified cell eQTL datasets

In order to validate the CT eQTLs defined by decon-eQTL, we utilized the eQTLs identified from purified neutrophils, CD4+ T-cells and CD14+ monocytes^9^. We first compared the absolute effect sizes from purified cells between significantly deconvoluted CT eQTLs, with non-significant deconvoluted CT eQTLs for this CT. For all three cell populations, effect sizes in our deconvoluted CT eQTLs were significantly higher compared to the effect size of eQTLs without a significant CT eQTL (Wlcoxon test, p-value ≤ 0.05, Fig. 4C). Next, we assessed the specificity of our deconvoluted CT eQTLs by evaluating CT-eQTL effect sizes in non-relevant cell subpopulations. For example, we compared the effect sizes of deconvoluted granulocyte eQTLs against those with non-significant deconvoluted granulocyte eQTLs using the effect sizes of purified CD4+ T-cell eQTLs. Notably, we observed no statistically significant differences using effect sizes from non-relevant cell subpopulations (see off-diagonal comparisons in Supplementary Fig. 8), further supporting the biological significance of our deconvoluted CT eQTLs.

To further demonstrate that the cell type eQTLs assigned by Decon-eQTL are biologically relevant, we have made use of the K27AC and K4ME1 epigenetic QTLs characterized using purified neutrophils, CD4+ T-cells and monocytes CD14+^9^. In a similar fashion as the above comparison of effect sizes with purified eQTLs, we compared the absolute effect sizes from both K27AC and K4ME1 QTLs from eQTLs for which Decon-eQTL detects a CT effect against the rest of whole blood eQTLs. We observed that for corresponding cell types, e.g. evaluating granulocyte CT eQTLs in K27AC QTLs from purified Neutrophils, the distribution of the absolute effect sizes is significantly higher for the chromatin mark QTLs (cmQTLs) than those non-significant CT eQTLs, which provide an epigenetic evidence that our method is able to assign correctly the cell type eQTL effects, as shown in the diagonal comparisons in both K27AC QTLS (Supplementary Fig.9) and for K4ME1 QTLs (Supplementary Fig.10). Notably, we observed that for the non-relevant cell subpopulations only one comparison, i.e. granulocytes v.s. CD14+ monocytes, show a statistically significant higher effect sizes for K27AC QTLs and K4ME1 QTLs, although the difference in effect sizes is less pronounced as the ones observed with corresponding cell types. For the rest of the non-relevant comparisons in the off-diagonal of both Supplementary Fig.9 and Supplementary Fig. 10, there are no statistically significant differences.

In addition to the comparison of effect sizes, we ascertained the allelic concordance between deconvoluted eQTLs and eQTLs from purified cell subtypes^9^. For each available CT (neutrophils, CD14+ monocytes, and CD4+ T cells), we evaluated whether the direction of the eQTL effect on deconvoluted CT eQTLs was the same as the one observed from purified cell subpopulations. Remarkably, the allelic concordance between the deconvoluted eQTLs and purified eQTLs was high across cell types: 99% for granulocyte eQTLs (compared to neutrophil eQTLs), 96% for CD14+ monocytes eQTLs, and 99% for CD4+ T cells (**Fig. 5A**). These rates of allelic concordance are significantly higher for deconvoluted granulocyte and CD4+ T-cell eQTLs compared to the those between whole blood eQTLs and eQTLs from purified cell subpopulations (**Fig. 5B**, Neutrophils, Fisher exact p-value = 3.91×10^6^, CD4+ T cells Fisher exact p-value = 0.005), whereas the allelic concordance for deconvoluted CD14+ monocyte eQTLs is the same as for whole blood eQTLs and purified CD14+ monocyte eQTLs (**Fig. 5B**). We also compared the allelic concordance of deconvoluted CT-eQTLs of a certain cell type against the eQTLs of non-relevant purified subpopulations. Interestingly, the allelic concordance across non-relevant cell subtypes is consistently lower (off-diagonal Supplementary Fig.11). The higher allelic concordance across CTs was seen between deconvoluted granulocyte eQTLs and CD14+ monocyte eQTLs with a 95% allelic concordance, which shows that the direction of effect is often shared between related cell types.

**Figure 5.**
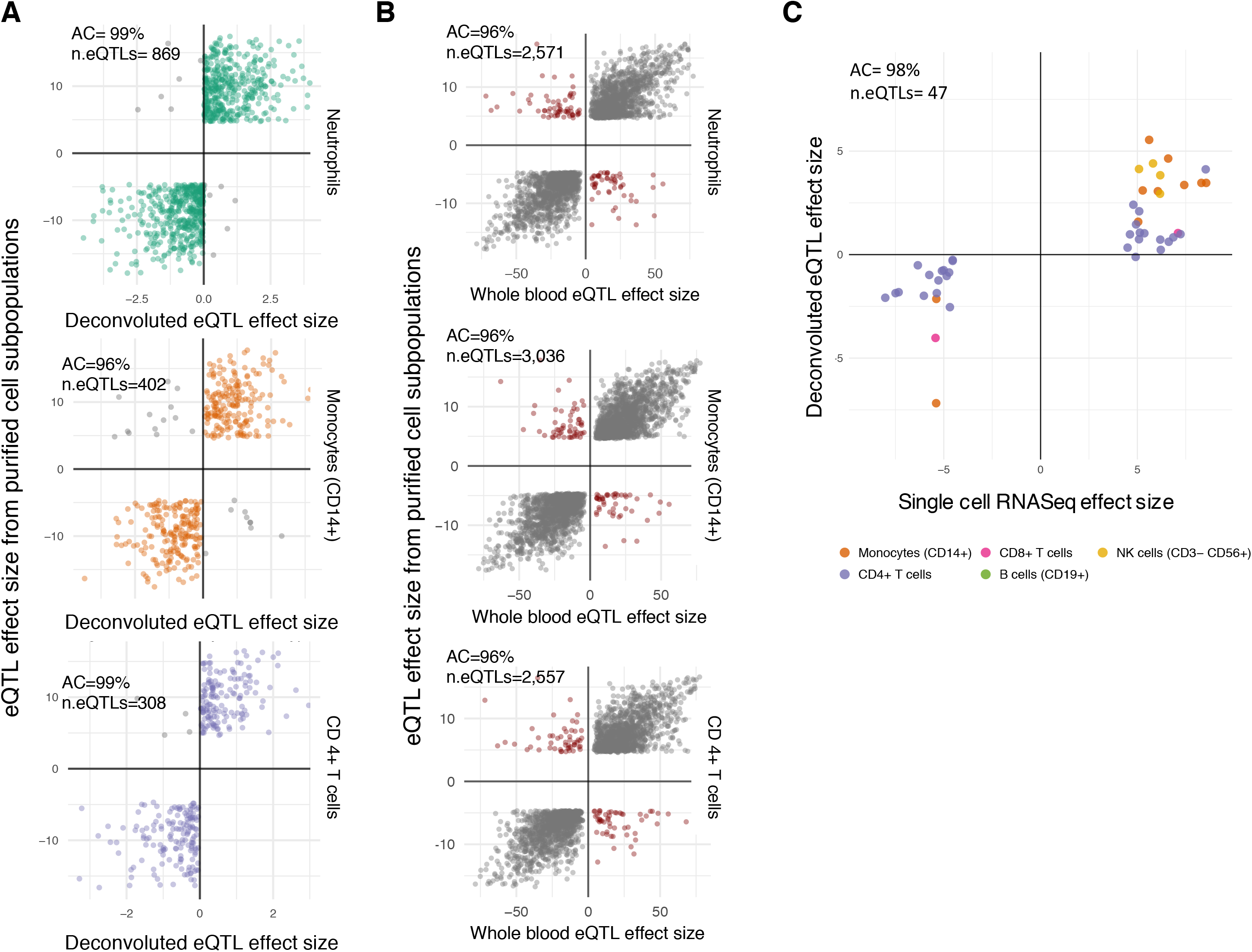
Allelic concordance of deconvoluted cell-type eQTLs with eQTLs from purified cells. Deconvoluted CT QTLs show high allelic concordance compared to eQTLs from purified cell subpopulations^9^. (A) for granulocyte eQTLs (orange), Decon-eQTL achieved an allelic concordance of 99% compared to eQTLs from purified neutrophils. Similarly, the allelic concordance were 96%and 99% for monocytes and CD4+ T cells, respectively. They are higher than those observed for whole blood eQTLs when comparing to eQTLs from purified subpopulations as shown in panel (B). Deconvoluted eQTLs show an allelic concordance of 95% for significant eQTLs obtained from single cell RNA-seq data ^24^ on monocytes CD14+, B cells, CD4+ T cells, CD8+ T cells and NK cells (C).

Finally, we evaluated the allelic concordance rates for CT eQTLs assigned by Decon-eQTL and K27AC QTLs from purified cell subpopulations, where we observed a consistently high allelic concordance rate: 92% for granulocyte eQTLs (in purified Neutrophils), 87% for CD14+ monocytes and 92% for CD4+ T cells (boxed diagonal comparisons in Supplementary Fig. 12). These concordance rates are significantly higher than the ones between the whole blood eQTLs and K27AC QTLs from purified cell subpopulations (Supplementary Fig 13) for neutrophils (Fisher exact test p-value = 9.06×10^−14^), CD14+ monocytes (Fisher exact test p-value = 3.33×10^−4^), C4+ T cells (Fisher exact test p-value = 8.64×10^−9^). Remarkably we also notice a consistent decrease on concordance rates when assessing the allelic concordance of CT eQTLs in K27AC QTLs of non-relevant cell subpopulations (off-diagonal compasons, Supplementary Fig. 12). Together, the results from allelic concordance rates between deconvoluted CT eQTLs and eQTLs/K27AC QTLs from purified cell subpopulations add a further layer of evidence supporting the biological relevance of deconvoluted CT eQTLs.

### CT eQTLs identified by Decon-eQTL in whole blood show allelic concordance with single-cell RNA-seq eQTLs

To replicate the deconvoluted CT eQTLs in the cell subtypes that were not available in Chen et al^9^. purified cell eQTLs, we utilized the recent single-cell RNA-seq eQTLs (sc-eQTLs) identified in CD14+ monocytes, NK cells, CD4+ T-cells, CD8+ T-cells, and B-cells^24^ We selected the top SNP per sc-eQTL pair for each of the cell types and compared it to the direction of the eQTL effect given by Decon-eQTL. Overall we observed an allelic concordance of 98% (Fig. 5C and Supplementary Table 3). This allelic concordance is higher than the one achieved on comparing the direction of whole blood eQTL effects with sc-eQTLs, where we observed an allelic concordance of 89% (Supplementary Fig. 14). Although the difference is not statistically significant (Fisher exact p-value = 0.1102), we expect that replication can be achieved for more rare cell types when single cell eQTL datasets with a larger sample size become available.

### Decon-QTL outperforms earlier methods

To our knowledge, our approach is the first to model the effect of multiple components of bulk blood RNA-seq to deconvolute cell type effects. Previous studies used an interaction effect between genotype and cell proportions of one specific cell type to detect the cell type eQTLs effects using whole blood gene expression^10,11^, or used the correlation of the eQTL effect with cell type proxy genes^10,11^.

The Westra *et al* method has often been used to detect cell type eQTL effects using bulk expression data and cell proportions^28–31^. In brief, it focuses on the effect of the GxE interaction (where E represents cell proportions) for explaining the variation in gene expression, and it only incorporates one cell type at each time. To properly compare Decon-eQTL with the Westra *et al* method, coined here ‘Westra method’, both methods were applied to the BIOS cohort, where we detected CT eQTLs for the six cell subpopulations. Replication of CT eQTLs from Westra method was done in the same way as described above for Decon-eQTL. We observed that the eGenes (i.e. genes with eQTLs) detected by the Westra method are significantly higher expressed for granulocytes (observed in purified neutrophils), CD4+ T cells and B cells, but not for CD14+ monocytes (Supplementary Fig. 15A). Next, we found that the distribution of effect sizes in eQTLs from purified cells is significantly higher for the CT eQTLs detected using the Westra *et al* method when compared to the rest of the whole blood eQTLs (boxed-diagonal comparisons in Supplementary Fig. 15B), showing similar results as the ones from Decon-eQTL. However, their performance differentiates when comparing effect sizes of eQTLs of non-relevant cell subpopulations (off diagonal comparisons in Supplementary Fig. 16), where the Westra method shows less CT specificity, mainly across neutrophils and CD14+ monocytes, as observed by a significant difference (Wilcoxon test p-value = 4×10^−08^, Fig S15B), whereas from Decon-eQTL this comparison yields a non significant difference (Wilcoxon test p-value = 5.2×10^−02^). This difference in effect sizes by the Westra method in non-relevant cell subpopulations is also observed for eQTLs detected in CD14+ monocytes by the Westra method when compared to CD4+ T cell effect sizes. These results suggest that the results obtained with the Westra method are not as specific as the ones detected by Decon-eQTL.

When comparing the allelic concordance rates between the direction of effects given by the interaction term from the Westra method and those found in eQTLs from purified cell subpopulations, we observed that the allelic concordance for granulocytes eQTLs, 99%, (evaluated in neutrophils) and for CD4+ T cells, 100% (Supplementary Fig. 16) is comparable as to those observed for Decon-eQTL (Fig.4A). Conversely, the allelic concordance rate for the CD14+ monocytes is only 28%, much lower than the results from Decon-eQTL(96%). Finally, for granulocytes, CD4+ T cell eQTLs and monocytes, we have overlapped the the results from Westra method and Decon-eQTL with the eQTLs from purified cell types (Chen et al) (Supplementary Fig. 17). For all three cell types, we found that Decon-eQTL is able to detect a larger number of eQTLs, with a similar replication rate as the Westra method.

The Zhernakova et al method^11^ uses modules of co-expressed genes from whole blood RNA-seq data to ascertain the effect on context/CT dependent eQTLs. We compared our Decon eQTL results with those from the Zhernakova method for neutrophils, CD4+ and CD8+ T-cells, CD14+ monocytes, and B-cells. The reported Z-scores for bulk whole blood eQTLs identified by Zhernakova *et al* were used to infer the allelic direction for each available CT. Again, we compared the direction of the eQTL effect with that of the purified neutrophils, CD4+ T-cell and CD14+ monocyte eQTLs^9^. Zhernakova et al. detected fewer CT eQTLs effects compared to Decon-eQTL(Fig. 3A for Decon-eQTL, Supplementary Fig. 18A). Although the eQTLs from the neutrophil module showed 100% concordance with the purified neutrophils, slightly outperforming Decon-eQTL (99% allelic concordance) (Supplementary Fig. 18B), the concordance rate for the other two cell types (80% for CD14+ monocytes module and 95% for CD4+ module) are lower than those from Decon-eQTL (96% and 99% respectively). Overall, these results demonstrate that Decon-eQTL is able to detect more CT eQTLs that can be replicated in purifiec eQTL dataset that previously reported methods, specially in not so abundant cell types such as CD14+ monocytes. However, the detection of interaction effects between genotype and cell proportions to dissect bulk (in this case whole blood) expression data and define cell type eQTLs remains an area of opportunity that could still be explored by the increase number of samples present in functional genomic cohorts and the greater number of purified eQTL dataset that can be used for validation.

## Discussion

We have developed a novel statistical framework, Decon2, which predicts the proportions of known cell subtypes using gene expression levels from bulk blood tissue (Decon-cell). Subsequently, these predicted cell proportions, together with genotype information and expression data, can be used to deconvolute a bulk eQTL effect into cell-type effects (Decon-eQTL). Using a set of samples with both whole blood RNA-seq data and cell frequencies of 73 cell subpopulations, we demonstrated that Decon-cell was able to predict 34 independent cell subpopulations. The performance of Deocn-cell has been extensively validated by using multiple independent cohorts and compared with existing methods. The obtained Decon-cell models were applied to a cohort of 3,189 samples with whole blood RNA-seq available, resulting in predicted cell counts for these samples. By integrating bulk expression data, genotype and predicted cell counts of BIOS cohort, Decon-eQTL was able to dissect whole blood eQTL effect into CT eQTLs without purifying immune cell subpopulations. Again the results of Decon-eQTL were validated by using several independent data types: 1) eQTLs from purified eQTL dataset, 2) chromatin QTLs purified eQTL dataset 3) gene expression from purified cell types. Compared with existing methods, Decon-eQTL consistently show superior performance. To sum up, the proposed framework is useful for analyze/re-analyzing both existing and new bulk bloodtissue datasets to detect cell-type eQTL effects, and can be applied and tested on other tissues once cell count proportions become available. This will improve our understanding of the functional role of SNPs associated to complex diseases, at the level of specific cell subtypes.

The main advantage of our method for predicting cell proportions by Decon-cell is that it does not rely on the gene expression measured in purified cell subtypes when defining signature gene sets. Moreover, our method does not require the definition of marker genes based on their differential expression compared to other cell subpopulations unlike previously reported methods^12^. The signature genes defined by Decon-cell are determined by a completely unsupervised approach using a regularized regression to select an optimal combination of genes to accurately predict a certain circulating cell proportion. Although the majority of these marker genes are differentially expressed across purified cell subpopulations, not all of them are. Nevertheless, these signature gene sets are still correlated to the cell proportions in whole blood. In summary, have shown that Decon-cell is able to accurately predict the proportions of circulating immune cell subpopulations in three independent cohorts and that within these cohorts it out-performs previously reported methods.

Our Decon-eQTL method for detecting an CT eQTL effect with bulk blood tissue expression data is, to our knowledge, the first attempt to simultaneously model bulk blood gene gene expression profiles into its major components. In contrast to a previous method, where single cell type (G x E) effects were evaluated one at a time^10,31^, Decon-eQTL incorporates all the major cell proportions simultaneously to better dissect the overall genetic effect of gene expression signal into cell subpopulations. We have validated our Decon-eQTL results by using eQTLs from purified neutrophils, CD14+ monocytes and CD4+ T-cells. Furthermore, we have shown that the eQTLs detected by Decon-eQTL have significantly higher effect sizes, specifically in the relevant cell subpopulations and they show an allelic concordance of at least 96%. Moreover, we have also shown the biological relevance of the deconvoluted CT eQTLs by validating our results on cmQTLs where CT eQTLs have significantly higher effect sizes and its allelic concordance rates are significantly higher than those of whole blood eQTLs. Finally, we have also demonstrated that Decon-eQTL can replicate CT eQTLs derived from single-cell RNA-seq data, showing a higher allelic concordance with sc-eQTLs compared to using only whole blood eQTL effects.

There are limitations in our method: the CT eQTLs detected by Decon-eQTL tend to be exclusive eQTL for the specific CT suggesting that the CT with the strongest eQTL effect was selected by Decon-eQTL. This is likely due to the partial collinearity present between CT proportions included in the model (as shown by their correlation structure in Supplementary Fig. 19A-B). Thus, the genetic effect of one cell type might be masked by another CT with correlated cell proportion. The highest correlation coefficient among cell types included in the model was 0.75 (between granulocytes and B cells). Therefore, a caveat to this is that by deconvoluting CT eQTLs for partially correlated cell proportions could lead to false negative results for the CTs with relatively weaker eQTL effects.

The proposed framework of Decon2 is generic for predicting cell subpopulations in bulk tissues (Decon-cell) and re-distribute the overall eQTL effect into cell types (Decon-eQTL). Both methods have been implemented into freely available software. In both R package and webtool, the models for predicting cell subpopulation in whole blood constructed and validated in this work are provided for people interested in estimating immune cell subpopulations in whole blood in health people with western european ethnicity, as our models were built using a Dutch cohort (500FG).

In summary, Decon2 is a computational method that can accurately assign CT effects in bulk blood eQTL datasets, which can be applied to any dataset for which genotypes and expression data is available to and potentially aid in our understanding of the molecular effects of genetic risk factors associated to complex diseases at cell type level. Our method makes it possible to create CT gene regulatory networks that could explain the different effects that each CT has on a complex disease in a cost-efficient way. Since Decon2 only requires gene expression and genotype information to deconvolute eQTLs, it is possible to re-analyze the existing bulk blood RNA-seq data for which genotypes are also available; this is where we would use Decon-cell to predict cell proportions in whole blood and obtain CT information on many more eQTLs from an increase in sample size. The methods behind Decon2 can be potentially generalized to use transcriptional profiles derived from any other type of bulk tissue in addition to whole blood, such as biopsies from tumors or other solid tissues implicated in complex disease etiology. Our methods can hence aid in the detection of genetic effects on gene expression in rare cell subpopulations in bulk tissues.

## Methods

### RNA-seq data collection in 500FG cohort

We selected a representative subset of 89 samples from the 500 participants of the 500FG cohort, which is part of the Human Functional Genomics Project (HFGP). Our subset was balanced for age and sex given the original distribution in the cohort, we performed RNA-seq in their whole blood samples. RNA was isolated from whole blood and subsequently globin transcripts were filtered by applying the Ambion GLOBINclear kit. The samples were then processed for sequencing using the library preparation kit lllumina TruSeq 2.0. Paired-end sequencing of 2×50-bp reads was performed on the lllumina HiSeq 2000 platform. The quality of the raw reads was checked using FastQC (http://www.bioinformatics.babraham.ac.uk/projects/fastqc/). Read alignment was performed with STAR 2.3.0^32,33^, using the human Ensembl GRCh37.75 as reference, whilst the aligned reads were sorted using SAMTools^34^. Lastly, gene level quantification of the reads was done using HTSeq^35^.

### RNA-seq preparation and data processing in the BIOS cohort

RNA was isolated from whole blood and subsequently globin transcripts were filtered by applying the Ambion GLOBINclear kit. Library preparation was performed using the lllumina TruSeq v2 library preparation kit. Next, lllumina HiSeq 2000 was used to performed paired-end sequencing of 2 × 50 bp reads while pooling 10 samples per lane and expecting >15 million read pairs per sample. By using CASAVA read sets were generated, retaining only reads that passed lllumina Chastity Filter for further processing.

Quality control of the reads was evaluated using FastQC (http://www.bioinformatics.babraham.ac.uk/projects/fastqc/). Adaptor sequences were trimmed out using cutadapt (v1.1) using default settings. Low quality ends of reads were removed using Sickle (v1.200) (https://github.com/najoshi/sickle).

Reads were then aligned using STAR 2.3.0e^33^. All SNPs present in the Genome of the Netherlands (GoNL) with MAF ≥ 0.01 were masked from the reads to avoid reference mapping bias. Read pairs with at most eight mismatches and mapping to at most five positions, were used. Quantification of counts per genes was done using Ensembl v.71 annotation (which corresponds to GENCODE v.16).

### Genotype data of the BIOS cohort

Genotype information was independently generated by each of the cohorts, further details on data collection and and methods used for genotyping can be found in their papers (CODAM^36^, LLDeep^16^, LLS^17^, RS^18^ and NTR^37^) Genotypes were harmonized to GoNL with Genotype Harmonizer^38^ and imputed using IMPUTE2^39^ using GoNL as reference panel. SNPs with an imputation score below 0.5, Hardy–Weinberg equilibrium P value smaller than 1×10^−4^, a call rate below 95% or a MAF smaller than 0.05 were filtered out. For further analysis only eSNPs from whole blood cis-eQTLs top effects were subsequently used in Decon-eQTL.

### Quantification of cell proportions in 500FG cohort

The inclusion criteria and further description of the participants of the 500FG cohort can be found at http://www.humanfunctionalgenomics.org. A total of 73 manually annotated immune cell subpopulations were quantified using 10-color flow cytometry. To minimize biological variability, cells were processed immediately after blood sampling and typically analyzed within 2–3 hr. Cell populations were gated manually as previously described^14^.

### *cis*-eQTLs in the BIOS cohort

For *cis*-QTL mapping, we tested association between genes and SNPs located within 250 kb of a gene center. SNPs with MAF ≥ 0.01, call rate = 1 and Hardy–Weinberg equilibrium p-value ≥ 0.0001 were included. eQTLs were declared to be significant at FDR < 0.05. Pre-processing of RNA-seq and QTL mapping was performed using a custom eQTL pipeline which has been previously described^11^.

### Prediction of cell proportions using gene expression levels from bulk tissue (Decon-cell)

We proposed that the abundance of molecular markers such as gene expression could be used as proxies to predict cell proportions. This can be represented as:

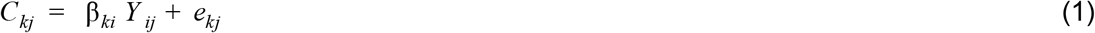

where expression data is Y_*ij*_ for genes *i* = 1, 2,…, G, and samples *j* = 1, 2, …, *N*, and cell count data is C_*kj*_ for sample j in cell type *k* (k = 1, 2, …, K), whilst β_*ki*_ represents the coefficients of gene *i* in determining cell counts of cell type *k* of a complex tissue and e_*kj*_ is the error term.

In order to select only the most informative genes for predicting cell counts, we implemented a feature selection scheme by applying an elastic net (EN) regularized regression^25^. In the EN algorithm, the β_*k*_ *Y* are estimated by minimizing:

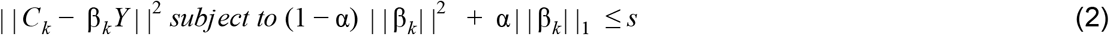

*s* is a tuning parameter that limits the number of features that will be included in the final predictor model. We estimate the best s per cell type by applying a 10-fold cross-validation approach, where the most optimal penalty parameter (α) was obtained.

### Normalization and correction of gene expression data for deconvolution of eQTL effects

Total read counts from HTSeq were first normalized using the trimmed means of M (TMM) values^32^. TMM expression values were log2 transformed. For predicting cell proportions, we used scaled expression data in both the 500FG and BIOS cohorts.

For the deconvolution of eQTLs, the expression was log2 transformed and corrected using a linear model for the effect of cohort, age, sex, GC content, RNA degradation rates, library size, and number of detected genes per sample. The corrected expression data is then exponentiated in order to maintain the original linear relationship across read counts (gene expression) and cell proportions.

### Deconvolution of eQTL effects (Decon-eQTL)

Decon-eQTL models the expression level in the bulk tissue by considering the genetic contribution of multiple cell types present in the system. For identifying the CT eQTL effect, the interaction term between a particular cell type and genotype was tested for statistically significance contribution to the explained variance on the expression levels of particular gene, while accounting for the remaining cell proportions.

If we consider a generic eQTL linear model for whole blood it can be described as:

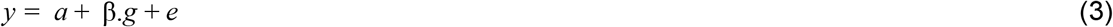

where *y* is the measured gene expression, *a* the modeled non-genetic dependent expression, *g* the genotype coded as 0, 1 or 2, β.*g* the genotype-dependent expression, and *e* the error, e.g. unknown environmental effects. Here all three terms are modeling the effect of the mixture of different cell types present in blood.

In an RNA-seq based gene expression quantification of a bulk tissue, one could express gene expression levels (*y*) as the sum of counts (ψ) per *K* cell types:

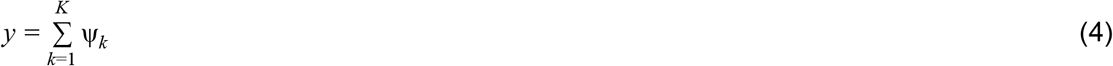

For every cell type the expression level has can be written as a generic eQTL model (equation 3) weighted by the cell proportions. ψ_*k*_ is a combination of the genetic and non genetic contribution of the cell type to *y*. The non-genetic contribution per cell type is β.*c* where *c* is the cell count proportions, while the genetic contribution is β_*k*_. *g* : *c_k_*. For *k* cell types the expression then is

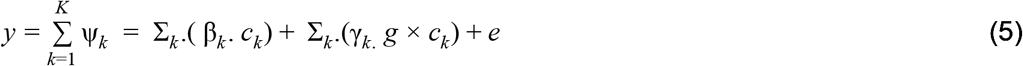

Where *y* is the measured expression levels, *k* is the total number of cell types, *c_k_* is the cell count proportions of cell type *k, g* is the genotype. And *e* is the error term. Since we are assuming a linear relationship between total gene expression and the levels of expression generated by each of the cell types composing a bulk tissue, the cell proportions are scaled to sum to 100%, such that the sum of the effect of the cell types equals the effect in whole blood. Here we assume that the true sum of the cell counts should be very close to 100% of the total PBMCs count, which is why we include the 6 cell types that together form the top hierarchy given the gating strategy used to quantify the cell subpopulations^14^. The genotype main effect is not include in the model as the sum of the genotype effect per cell type should approximate the main effect.

Because the contribution of each of the cell types to expression level *y* can not be negative, we constrain the terms of the model to be positive by using Non-Negative Least Squares^40,41^ to fit the parameters to the measured expression levels. However, if the allele that has a negative effect on gene expression is coded as 2, the best fit would have a negative interaction term, which would be set to 0. To address this we want the allele that causes a positive effect on gene expression to always be coded as 2. However, the effect of an allele has can be different per cell type, therefore the coding of the SNP should also be different per cell type. Therefore, we run the model multiple times, each time swapping the genotype encoding for one of the interaction terms. The encoding that gives the lowest R-squared is then chosen as the optimal genotype encoding. For the encoding we limit the amount of genotypes that have an opposite genotypic encoding to maximum of one interaction term, as we have observed that there no significant difference compared to using all possible configurations and this limits the amount of models that have to be run from k^2^to (2*k)+2.

To test if there is a CT interaction effect we run the linear model of equation 5. and, for each CT, run the same model with the cell proportion:genotype interaction term removed. E.g. when testing two cell types the full model is

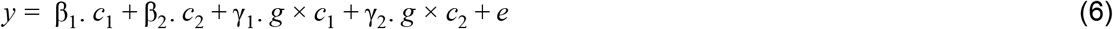

and the two models with the interaction terms removed are

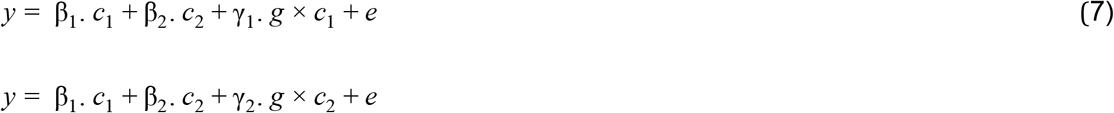

For both the full model and the CT models we calculated the sum of squares using the different genotype configurations detailed above. For both the full and the CT models we then selected the genotype configuration with lowest sum of squares. Then, for each CT, we test if full model can significantly explain more variance than the CT model using an ANOVA.

We have then applied our strategy to 16,362 significant whole blood cis-eQTLs top effects that were detected using the BIOS cohort. We then correct the p-values for multiple testing using FDR by each of the cell types, *e.i*. Granulocyte eQTL p-values were corrected for 16,362 tests, in the same way CD4+ T cells eQTL p-values were corrected for the exact same number of tests.

## Contributions

C.W., L.F. and YL initialized the study. Y.L. and L.F. directed and supervised the project. Y.L. developed the statistical framework, together with L.F. R.A-G, N.K., L.F., and Y.L., performed data analysis and interpretation. J.D.T. was involved in the initial analysis. N.K. and R.A-G. made the software and webtool. A.C, U.V., M. Z, X.C., O.B.B., Z.B., I.R.P., P.D., C.J.X., M.S., I.J., S.W., I.J., S.S., V.K., H.J.P.M.K., L.A.B.J., M.G.N. and C.W. contributed to data collection, data analysis and interpretation. R.A-G, N.K., L.F., and Y.L. draft and revised the manuscript. All authors have read and approved the manuscript.

## Acknowledgements

We thank K Mc Intyre and J Senior for editing the final text. We thank T. Spenkelink for the DeconCell web tool design. L.F. is supported by grants from the Dutch Research Council (ZonMW-VIDI 917.164.455 to M.S. and ZonMW-VIDI 917.14.374 to L.F.), and by an ERC Starting Grant, grant agreement 637640 (ImmRisk). Y.L. was supported by an ZonMW-OffRoad grant (91215206). The HFGP is supported by a European Research Council (ERC) Consolidator grant (ERC 310372). This study was further supported by an IN-CONTROL CVON grant (CVON2012-03) and a Netherlands Organization for Scientific Research (NWO) Spinoza prize (NWO SPI 94-212) to M.G.N.; an ERC advanced grant (FP/2007-2013/ERC grant 2012-322698) and an NWO Spinoza prize (NWO SPI 92-266) to C.W.; a European Union Seventh Framework Programme grant (EU FP7) TANDEM project (HEALTH-F3-2012-305279) to C.W. and V.K.;. RJX was supported by National Institutes of Health (NIH) grants - DK43351, AT009708, Al137325. A CONACYT-I2T2 scholarship (382117) to R.A-G. The Biobank-Based Integrative Omics Studies (BIOS) Consortium is funded by BBMRI-NL, a research infrastructure financed by the Dutch government (NWO 184.021.007). We thank the UMCG Genomics Coordination center, the UG Center for Information Technology and their sponsors BBMRI-NL & TarGet for storage and compute infrastructure.

## Supplementary Materials

### Supplementary figures

**Supplementary Figure 1: Prediction performance of Decon-cell within 500FG:** The Y-axis represents the 73 immune cell types quantified by FACS in the 500FG cohort. The bar plot on the left panel shows the mean Prediction Performance (Spearman correlation coefficient between predicted and measured cells across 100-fold cross validations). On the right panel, box plots represent the distribution of the Prediction Performance within 100 iterations of the cross validations. A cutoff of mean Prediction Performance ?0.5 was applied to define predictable cell types (green).

**Supplementary Figure 2. Signature genes selected for prediction of cell proportions by Decon-cell: (A)** Total number of marker genes (genes selected in ? 80% of all models in the 100 iterations) per predictable cell type. Different colors indicate different subpopulations. **(B)** The number of genes significantly correlated with cell counts (Spearman correlation, adjusted P ≤ 0.05) (y-axis) shows the total number of significantly correlated genes, while the x-axis shows the prediction performance (x-axis). **(C)** Distributions of the total number of “strongly” correlated genes (absolute Spearman correlation ≥ 0.3) between predictable and unpredictable cell subpopulations.

**Supplementary Figure** 3. **Comparison of prediction performance between Decon-cell and other existing methods.** (A) Performance of Decon-cell: he measured (x axis) and predicted cell proportions (y-axis) were compared for neutrophils (given by granulocytes in 500FG), lymphocytes and monocytes CD14+ and granulocytes three independent cohorts (shown by row, from top to bottom: LLDeep (n= 627), LLS (n= 660), RS (n= 773)). (B) Comparison of prediction performance for Decon-cell, CIBERSORT and xCell in three independent cohorts for a total of 4 major immune subpopulations.

**Supplementary Figure 4. Prediction performance of xCell and CIBERSORT in three independent Dutch populations** (LLDeep, n= 627; LLS, n= 660; RS, n= 773). (A) Scatter plots showing on the x-axis the measured cell proportions of circulating immune cells and the xCell enrichment score on the y-axis. (B) Scatter plots showing on the x-axis the measured cell proportions of circulating immune cells and the predicted cell proportions given by CIBERSORT)

**Supplementary Figure 5. Expression of marker genes selected by Decon-cell.** Expression levels (scaled, log2(TPM+1) of signature genes in the data in three purified cell subpopulations: CD4+ T cells **(A),** neutrophils/granulocytes **(B)** and monocytes **(C)** in the data from the BLUEPRINT. Cell subpopulations are indicated in different colors by columns. Correlation of each of the signature genes and the cell subpopulation percentage in 500FG cohort is shown on green bar at the left-hand side of heatmaps figure,i.e. darker green correspond to higher correlations.

**Supplementary Figure 6. Many of the deconvoluted eQTL are cell type exclusive.** The colored bar plot on the left shows the total number of significantly deconvoluted eQTLs in whole blood eQTLs (as shown also in Figure 2A). The gray bar plot shows the total number of eQTLs shared across the possible combinations of the six cell subpopulations under study.

**Supplementary Figure 7. Variation of gene expression across samples for deconvoluted cell-type eQTLs genes in whole blood.** Granulocyte eQTL genes show significantly higher variance across the BIOS samples (F test p-value ≤ 0.05) compared to those from monocytes, CD4+ T cells, CD8+ T cells, B cells and NK cells.

**Supplementary Figure 8. Validation of deconvoluted eQTLs using effect sizes of eQTLs from purified cells.** Deconvoluted eQTLs (FDR ≤ 0.05) from BIOS cohort show a significantly bigger effect size in purified cell eQTLs^9^ from their relevant cell subtype compared to other whole blood eQTLs (diagonal boxed comparisons). The off-diagonal comparisons show that these eQTL genes are specific to a cell subpopulation because the differences in effect sizes are non-significant in all but one (CD4+ T cell eQTL genes in monocyte-derived eQTLs).

**Supplementary Figure 9. Validation of deconvoluted eQTLs using effect sizes of** K27AC **QTLs from purified cells.** Deconvoluted eQTLs (FDR ≤ 0.05) show a significantly bigger effect size for K27AC QTLs which have peaks located in the promoter region of the the eGenes from their relevant cell subtype compared to the rest of the significant whole blood eQTLs (diagonal boxed comparisons). The off-diagonal comparisons show that these eQTL genes are specific to a cell subtype because the differences in effect sizes are non-significant in all but the comparisons across Neutrophils and Monocytes (CD14+).

**Supplementary Figure 10. Validation of deconvoluted eQTLs using effect sizes of** K4ME1 **QTLs from purified cells.** Deconvoluted eQTLs (FDR ≤ 0.05) show a significantly bigger effect size for K4ME1 QTLs (where the eGenes is the closest gene tagging the K4ME1 QTLs peak)from their relevant cell subtype compared to the rest of the significant whole blood eQTLs (diagonal boxed comparisons). The off-diagonal comparisons show that these eQTL genes are specific to a cell subtype because the differences in effect sizes are non-significant in all but the comparisons between neutrophils and monocytes (CD14+).

**Supplementary Figure 11. Validation of deconvoluted eQTLs using allelic concordance with eQTLs results from purified cells.** Deconvoluted eQTLs (FDR ≤ 0.05) show a high allelic concordance in their respective purified cell eQTLs. Top row shows allelic concordance of deconvoluted granulocyte eQTLs (all in green) against neutrophils, monocytes and CD4+ T cells. Second row shows deconvoluted monocyte eQTLs against purified cell eQTLs in the same order as top row; bottom row shows the same comparisons as for deconvoluted CD4+ eQTLs. Allelic concordance of the off-diagonal (comparing deconvoluted eQLTs with non-relevant cell types) show a consistent decrease in allelic concordance.

**Supplementary Figure 12. Validation of deconvoluted eQTLs using allelic concordance with K27AC results from purified cells.** Deconvoluted eQTLs (FDR ≤ 0.05) show a high allelic concordance in their respective purified cell K27AC QTLs. Top row shows allelic concordance of deconvoluted granulocyte eQTLs (all in green) against neutrophils, monocytes and CD4+ T cells derived K27AC QTLs. Second row shows deconvoluted monocyte eQTLs (all in orange) against purified cell K27AC QTLs in the same order as top row; bottom row shows the same comparisons as for deconvoluted CD4+ eQTLs (all in purple). Allelic concordance of the off-diagonal (comparing deconvoluted eQLTs with non-relevant cell types) show a consistent decrease in allelic concordance when compared to the relevant cell type comparisons.

**Supplementary Figure 13. Allelic concordance between whole blood eQTLs and K27AC QTLs** for purified neutrophils, CD14+ monocytes and CD4+ T cells.

**Supplementary Figure 14. Comparison of whole blood eQTLs with eQTLs from single cell RNA-seq** Whole blood eQTLs show 89% allelic concordance for significant eQTLs derived from single-cell RNA-seq data, comprising monocytes CD14+, B cells, CD4+ T cells, CD8+ T cells and NK cells.

**Supplementary Figure 15. Validation of cell type eQTLs detected in the BIOS cohort using Westra** *et at*, **method:** (A) Expression of eGenes in purified cell subpopulations from BLUEPRINT (green for granulocyte eQTL genes showing expression for purified neutrophils; orange for monocytes; purple for CD4+ T cells; pink for B cells). (B) CT eQTLs detected by the Westra method show a significantly larger effect size in purified cell eQTLs^11^ compared to the rest of the whole blood eQTLs. Boxed-diagonal show the comparisons with relevant cell types, were the effect differences are stronger.

**Supplementary Figure 16. Allelic concordance rates of cell type eQTLs detected using the Westra** *et al* **method and eQTLs from purified cells.** Top row shows allelic concordance of granulocyte CT eQTLs against neutrophils, monocytes and CD4+ T cells. Second row shows CT monocyte eQTLs against purified cell eQTLs in the same order as top row; bottom row shows the same comparisons for CT CD4+ eQTLs.

**Supplementary Figure 17. Comparison of Decon-eQTL with Westra** *et al* **method.** Overlap of CT eQTLS detected with Decon-eQTL, the Westra *et al* method and those found to be significant in purified cell subpopulations, for granulocyte QTLs (A), CD4+ T cells (B), and monocytes (C).

**Supplementary Figure 18. Comparison of Decon-eQTL with other methods for detecting cell type eQTLs.** Total number of eQTLs per cell proportion module obtained by Zhernakova et al. (Nat Gen, 2017) (A). Allelic concordance between overall z-score for eQTLs from neutrophil, monocytes and CD4+ T cell modules against the effect size of purified eQTLs from neutrophils, monocytes and CD4+ T cells.

**Supplementary Figure 19. Distribution and correlation among circulating cell proportions.** (A) With 89 samples from 500FG, the scatter plots show the correlations between different cell subpopulations. Blue line indicates a fitted linear model. Diagonal plots depict the overall density distribution per cell type. Upper right triangle shows the Pearson correlation coefficient for each pairwise comparison. (B) shows correlations between different cell subpopulations in the BIOS cohort, which were obtained by prediction using Decon-cell.

### Supplementary Tables

**Supplementary table 1**: Ensembl IDs and symbol names of the marker genes selected by Decon-cell for the 34 predictable circulating immune cell proportions.

**Supplementary table** 2: Summary statistics from Decon-eQTLs for the 16,362 whole blood eQTLs.

