## Supplementary Figures for "Deconvolution of bulk blood eQTL effects into immune cell subpopulations"

### Supp. Figure 1.

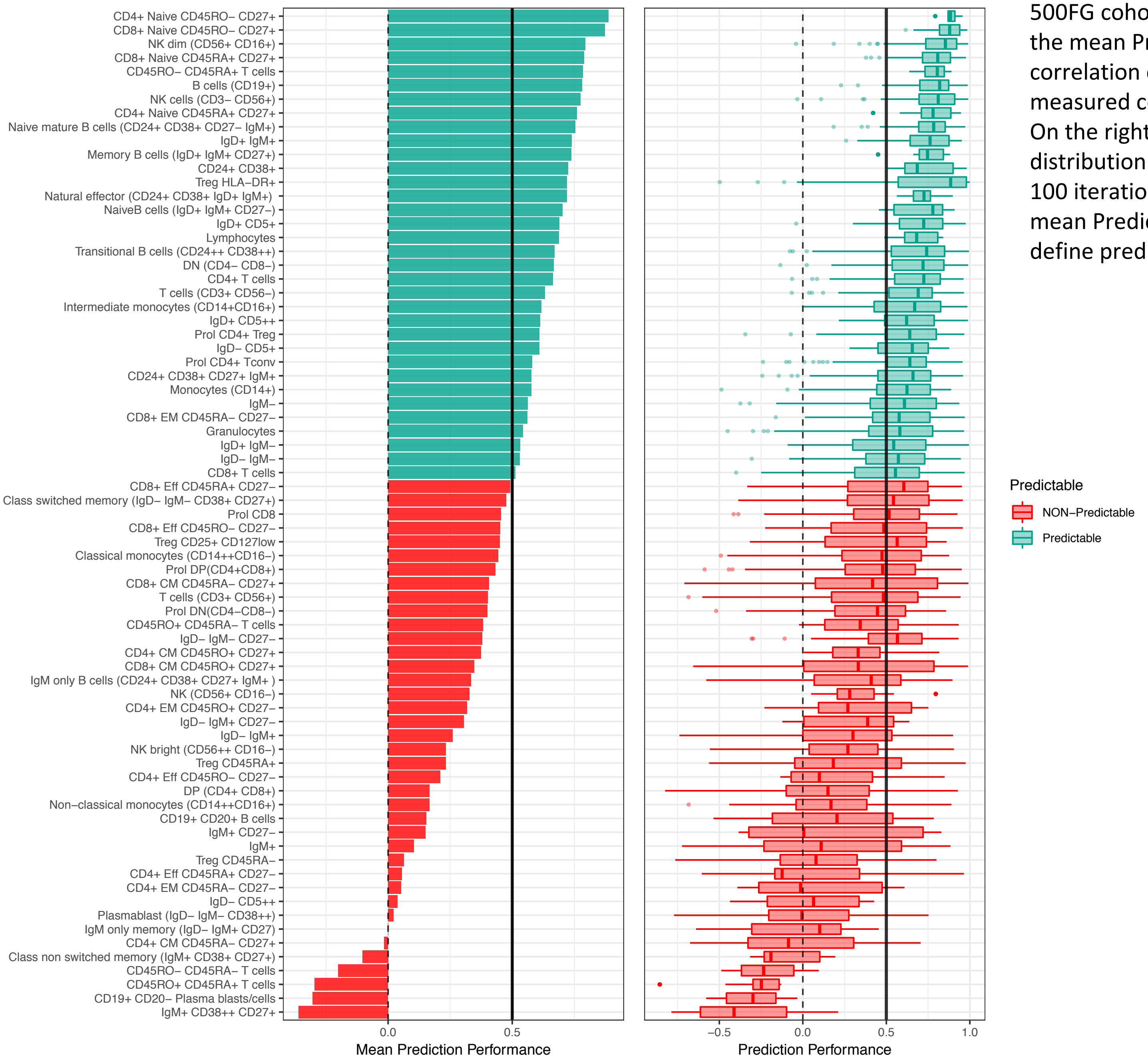


**Supplementary Figure 1: Prediction performance of Decon-cell within 500FG**: The Y-axis represents the 73 immune cell types quantified by FACS in the 500FG cohort. The  bar plot on the left panel shows the mean Prediction Performance (Spearman correlation coefficient between predicted and measured cells across 100-fold cross validations). On the right panel, box plots represent the distribution of the Prediction Performance within 100 iterations of the cross validations. A cutoff of mean Prediction Performance ≥0.5 was applied to define predictable cell types (green).

### Supp. Figure 2.

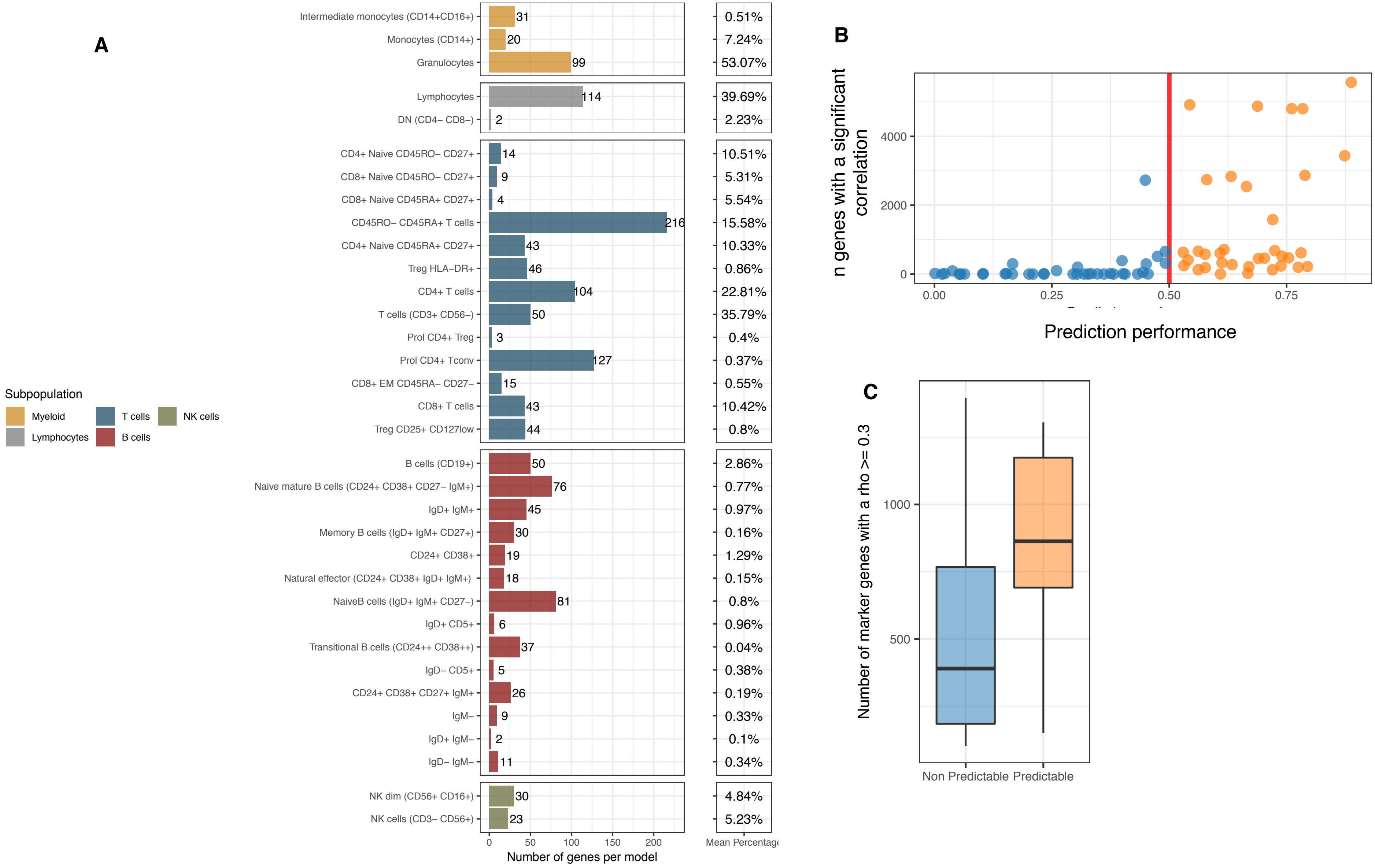


**Supplementary Figure 2. Signature genes selected for prediction of cell proportions by Decon-cell: (A)** Total number of marker genes (genes selected in ≥ 80% of all models in the 100 iterations) per predictable cell type. Different colors indicate different subpopulations. **(B)** The number of genes significantly correlated with cell counts (Spearman correlation, adjusted P ≤ 0.05)  (y-axis) shows the total number of significantly correlated genes , while the x-axis shows the prediction performance (x-axis). **(C)** Distributions of the total number of “strongly” correlated genes (absolute Spearman correlation ≥ 0.3) between predictable and unpredictable cell subpopulations.

Supp. Figure 3.

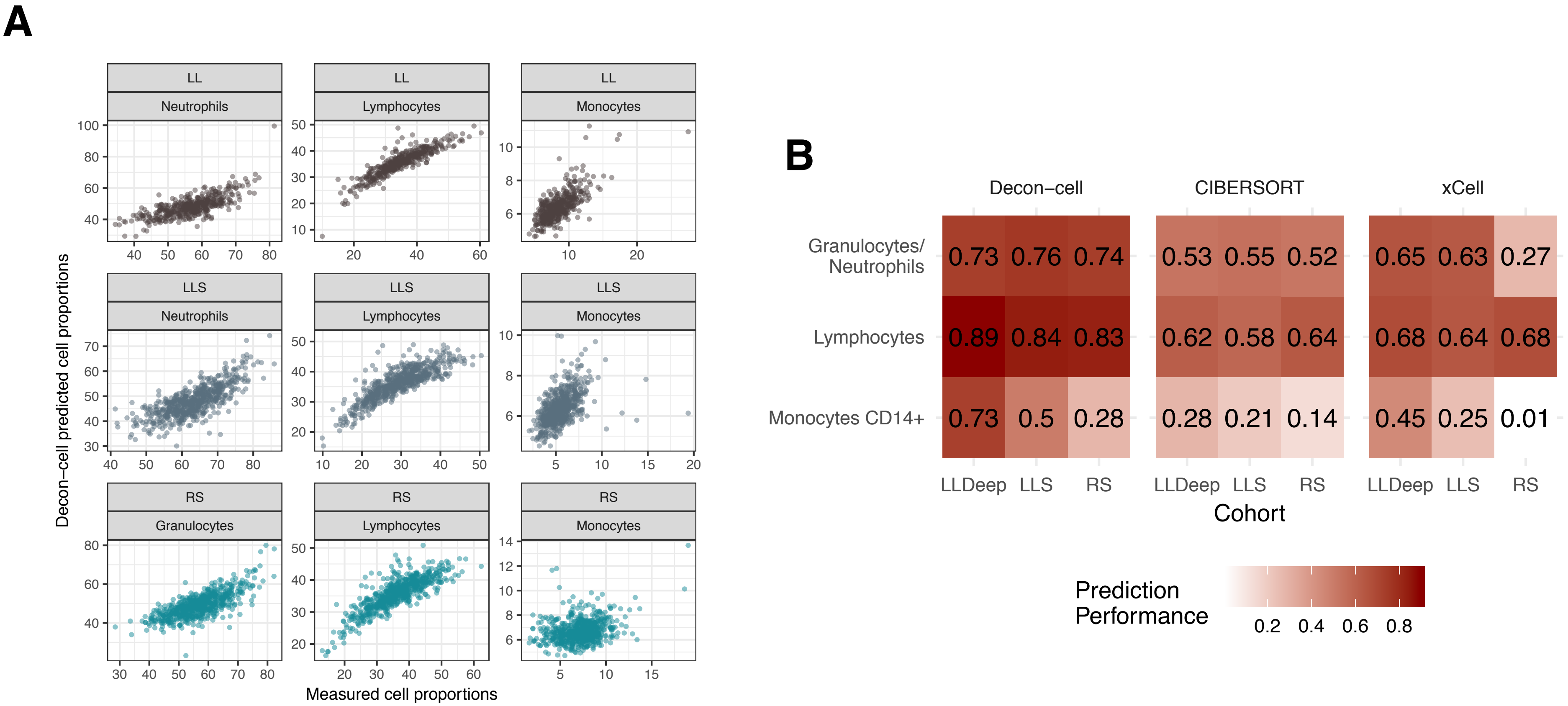


**Supplementary Figure 3. Comparison of prediction performance between Decon-cell and other existing methods.** (A) Performance of Decon-cell: he measured (x axis) and predicted cell proportions (y-axis) were compared for neutrophils (given by granulocytes in 500FG), lymphocytes and monocytes CD14+ and granulocytes three independent cohorts (shown by row, from top to bottom: LLDeep (n= 627 ), LLS (n= 660) , RS (n= 773)). (B) Comparison of prediction performance for Decon-cell, CIBERSORT and xCell in three independent cohorts for a total of 4 major immune subpopulations.

Supp. Figure 4.

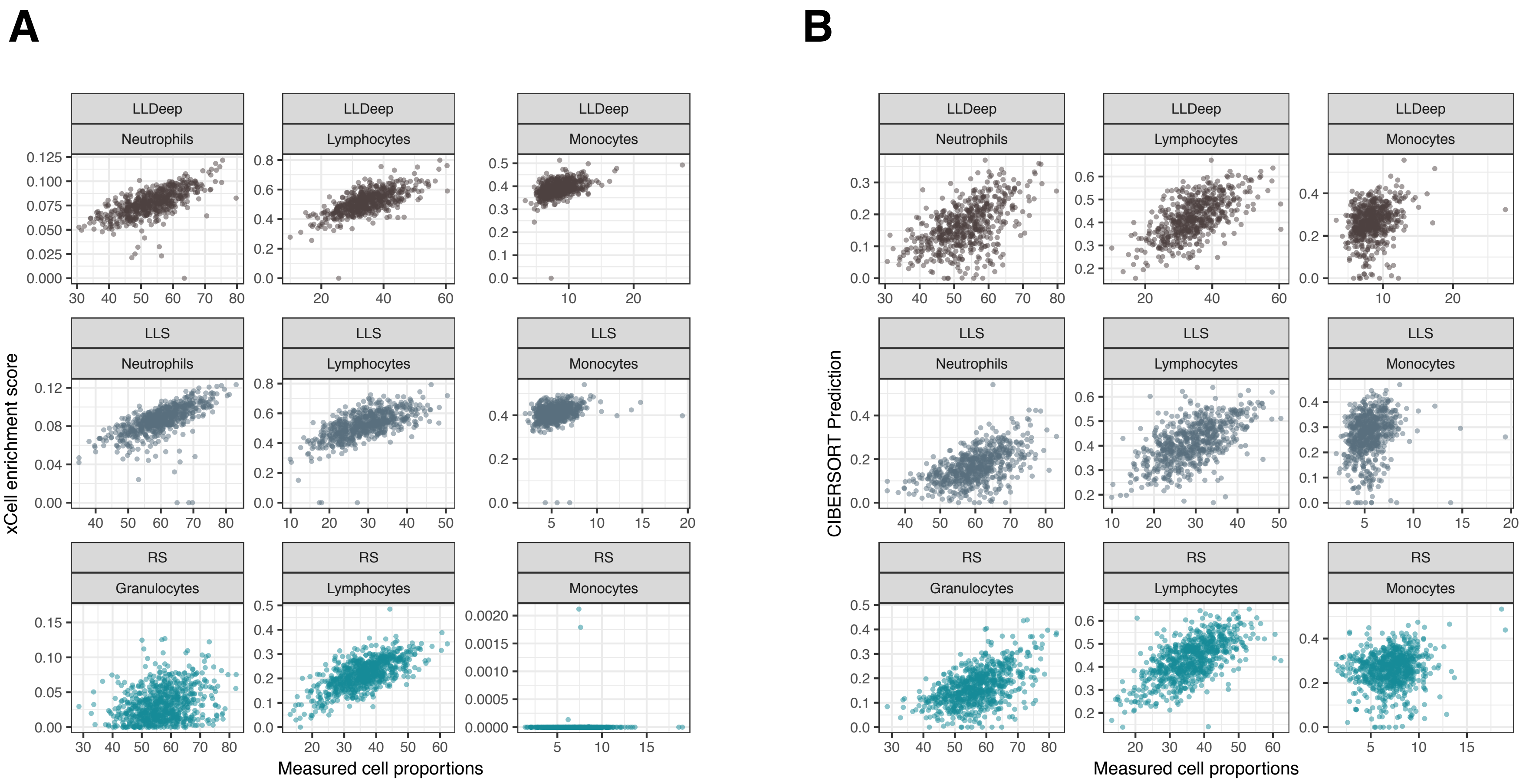


**Supplementary Figure 4. Prediction performance of xCell and CIBERSORT in three independent Dutch populations (**LLDeep, n= 627; LLS, n= 660; RS, n= 773)**.** (A) Scatter plots showing on the x-axis the measured cell proportions of circulating immune cells and the xCell enrichment score on the y-axis. (B) Scatter plots showing on the x-axis the measured cell proportions of circulating immune cells and the predicted cell proportions given by CIBERSORT)

Supp.Figure 5.

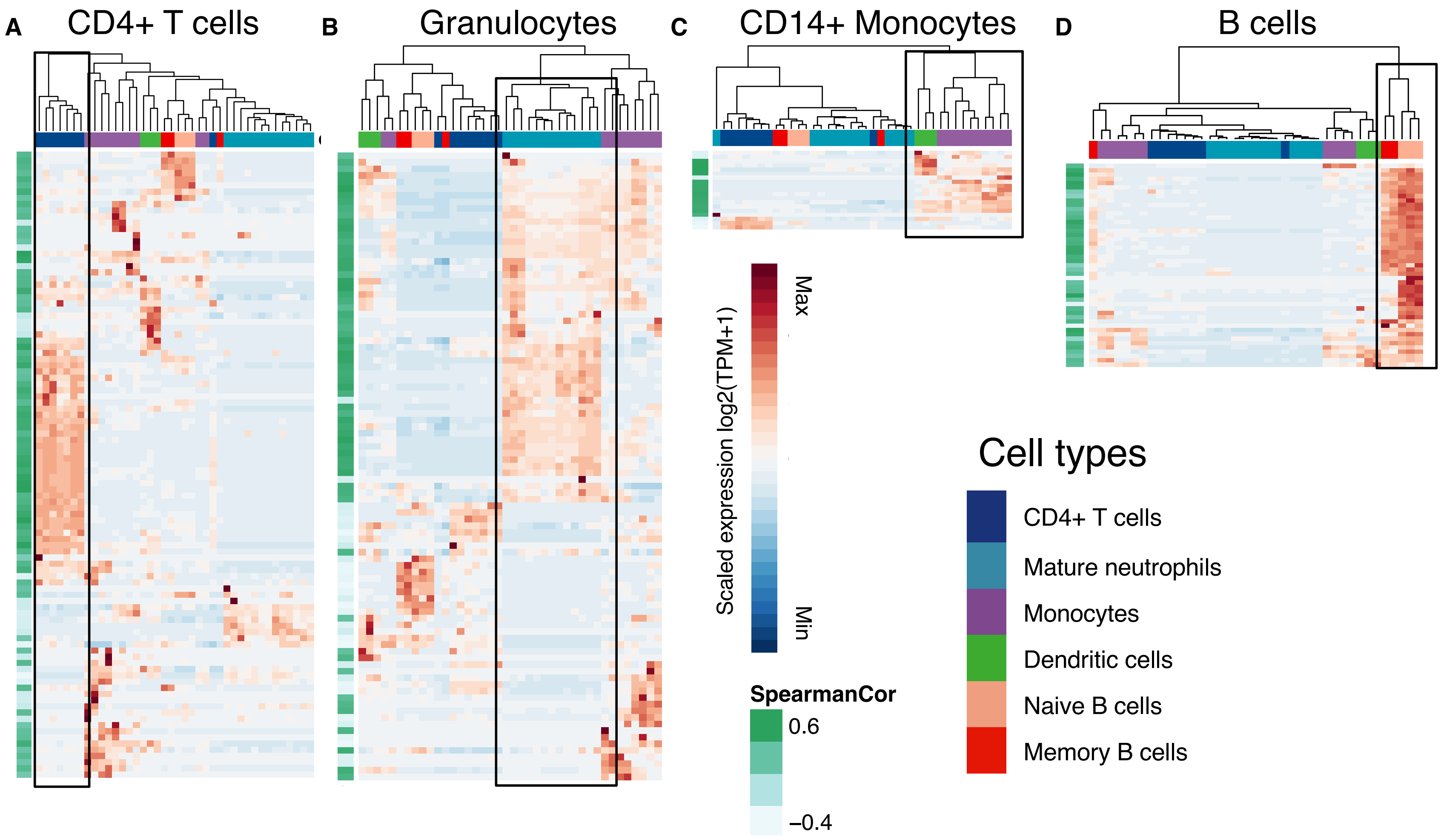


**Supplementary Figure 5. Expression of marker genes selected by Decon-cell.** Expression levels (scaled, log2(TPM+1) of signature genes in the data in three purified cell subpopulations: CD4+ T cells **(A)**, neutrophils/granulocytes **(B)** and monocytes **(C)** in the data from  the BLUEPRINT. Cell subpopulations are indicated in different colors by columns. Correlation of each of the signature genes and the cell subpopulation percentage in 500FG cohort is shown on green bar at the left-hand side of heatmaps figure,i.e. darker green correspond to higher correlations.

Supp.Figure 6.

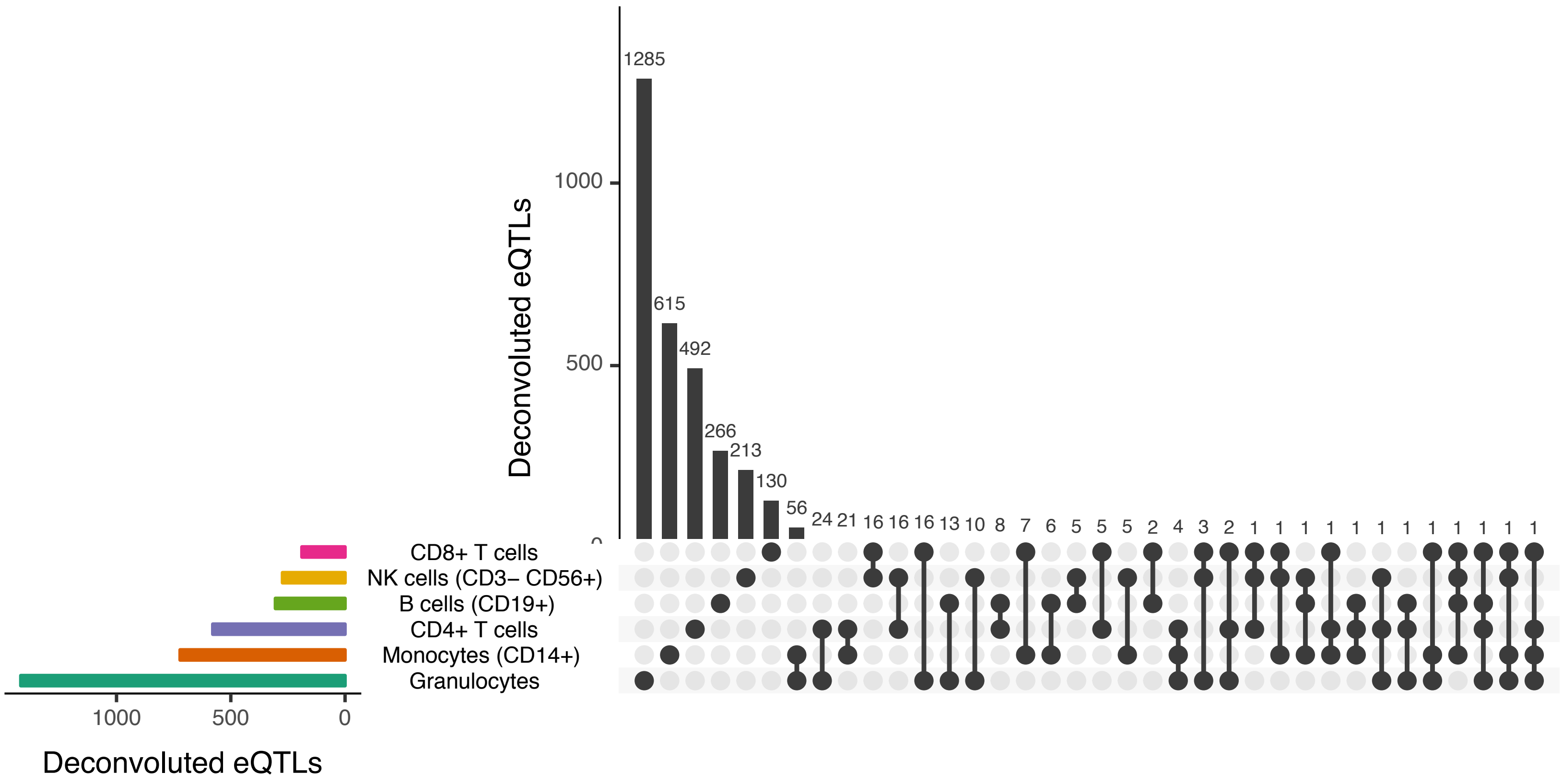


**Supplementary Figure 6. Many of the deconvoluted eQTL are cell type exclusive.** The colored bar plot on the left shows the total number of significantly deconvoluted eQTLs in whole blood eQTLs (as shown also in Figure 2A). The gray bar plot shows the total number of eQTLs shared across the possible combinations of the six cell subpopulations under study.

Supp.Figure 7

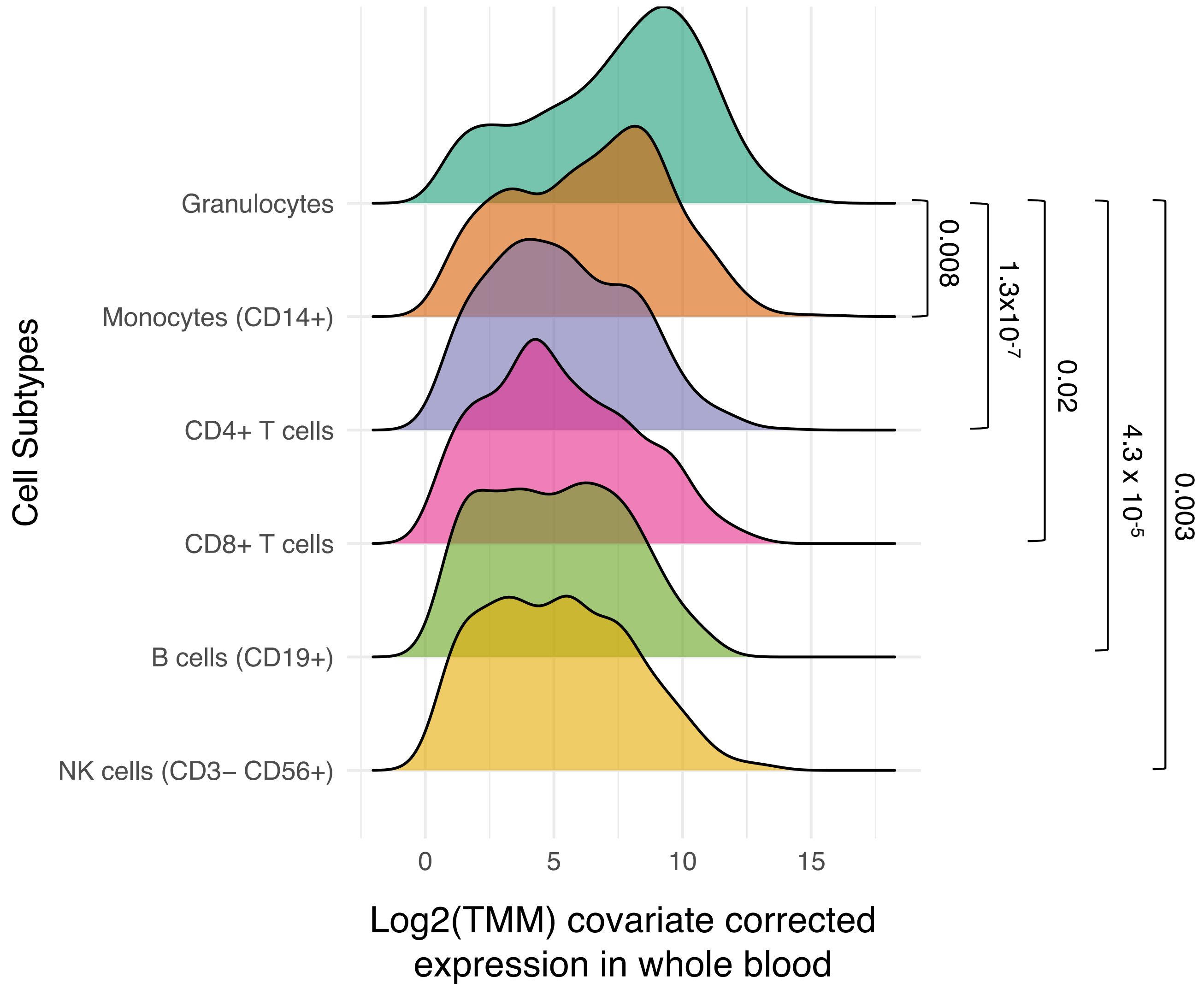


**Supplementary Figure 7. Variation of gene expression across samples for deconvoluted cell-type eQTLs genes in whole blood.** Granulocyte eQTL genes show significantly higher variance across the BIOS samples (F test p-value ≤ 0.05) compared to those from monocytes, CD4+ T cells, CD8+ T cells, B cells and NK cells.

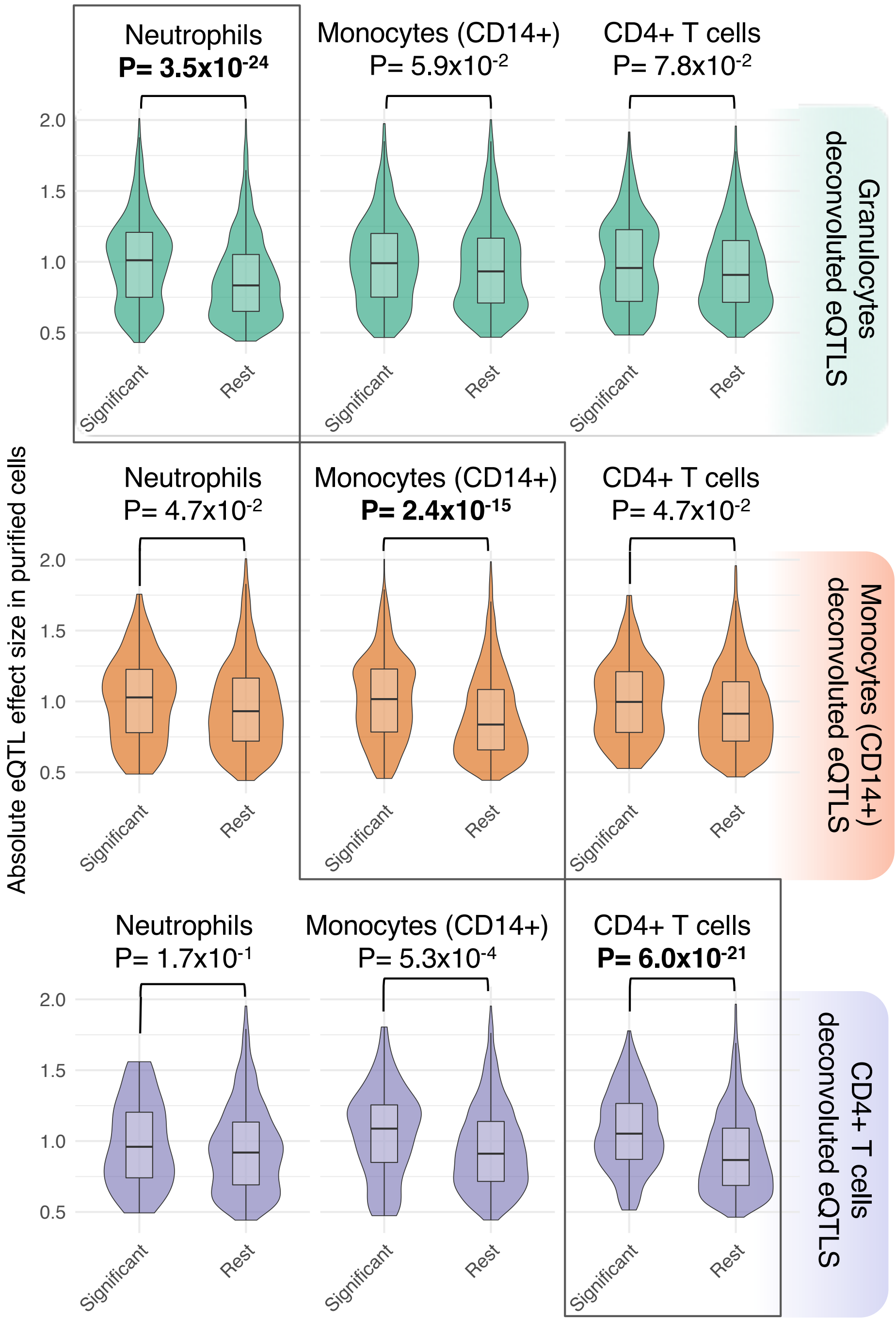


**Supplementary Figure 8. Validation of deconvoluted eQTLs using effect sizes of eQTLs from purified cells.** Deconvoluted eQTLs (FDR ≤ 0.05) from BIOS cohort show a significantly bigger effect size in purified cell eQTLs[9] from their relevant cell subtype compared to other whole blood eQTLs (diagonal boxed comparisons). The off-diagonal comparisons show that these eQTL genes are specific to a cell subpopulation because the differences in effect sizes are non-significant in all but one (CD4+ T cell eQTL genes in monocyte-derived eQTLs).

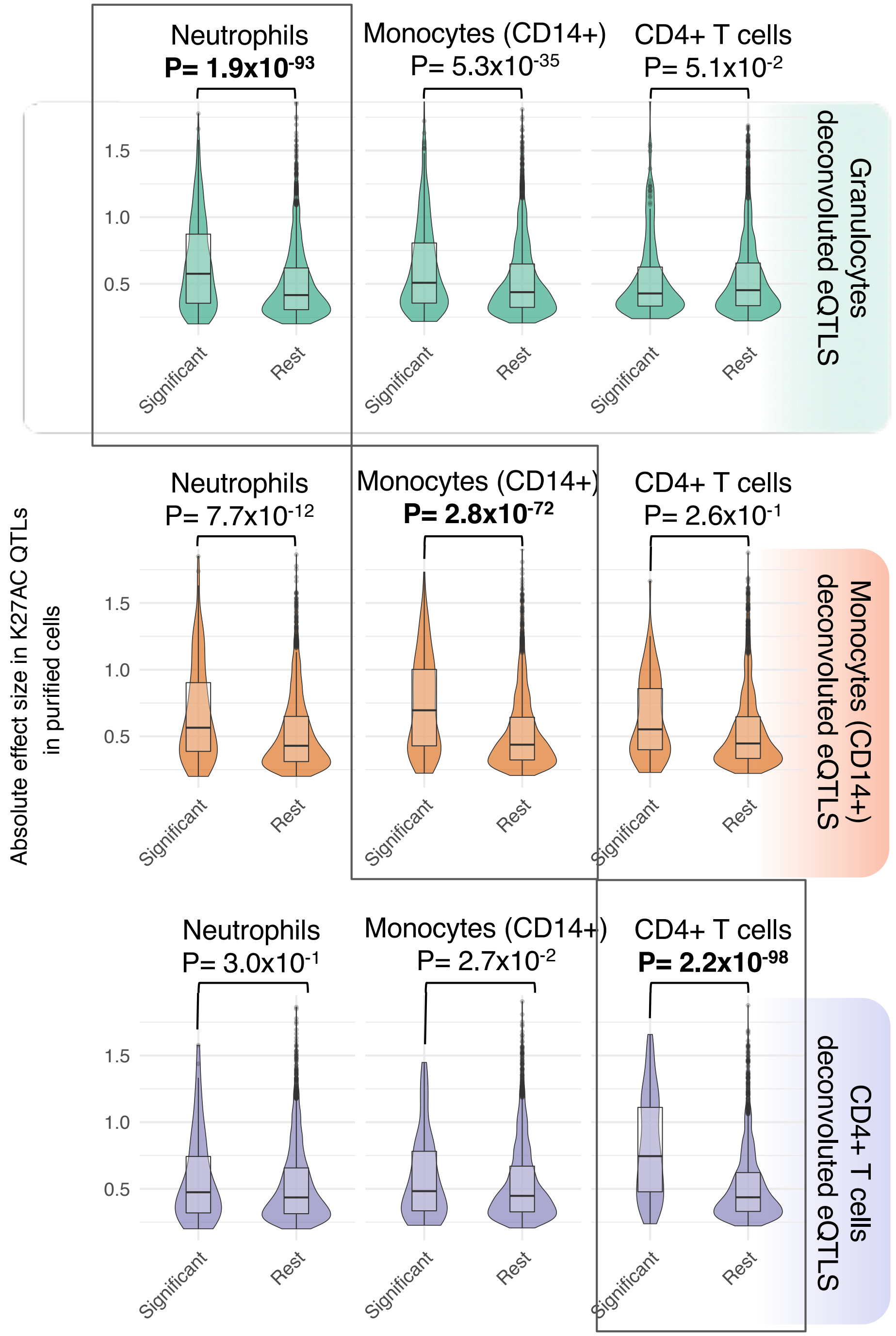


**Supplementary Figure 9. Validation of deconvoluted eQTLs using effect sizes of** K27AC **QTLs from purified cells.** Deconvoluted eQTLs (FDR ≤ 0.05) show a significantly bigger effect size for K27AC QTLs which have peaks located in the promoter region of the the eGenes from their relevant cell subtype compared to the rest of the significant whole blood eQTLs (diagonal boxed comparisons). The off-diagonal comparisons show that these eQTL genes are specific to a cell subtype because the differences in effect sizes are non-significant in all but the comparisons across Neutrophils and Monocytes (CD14+).

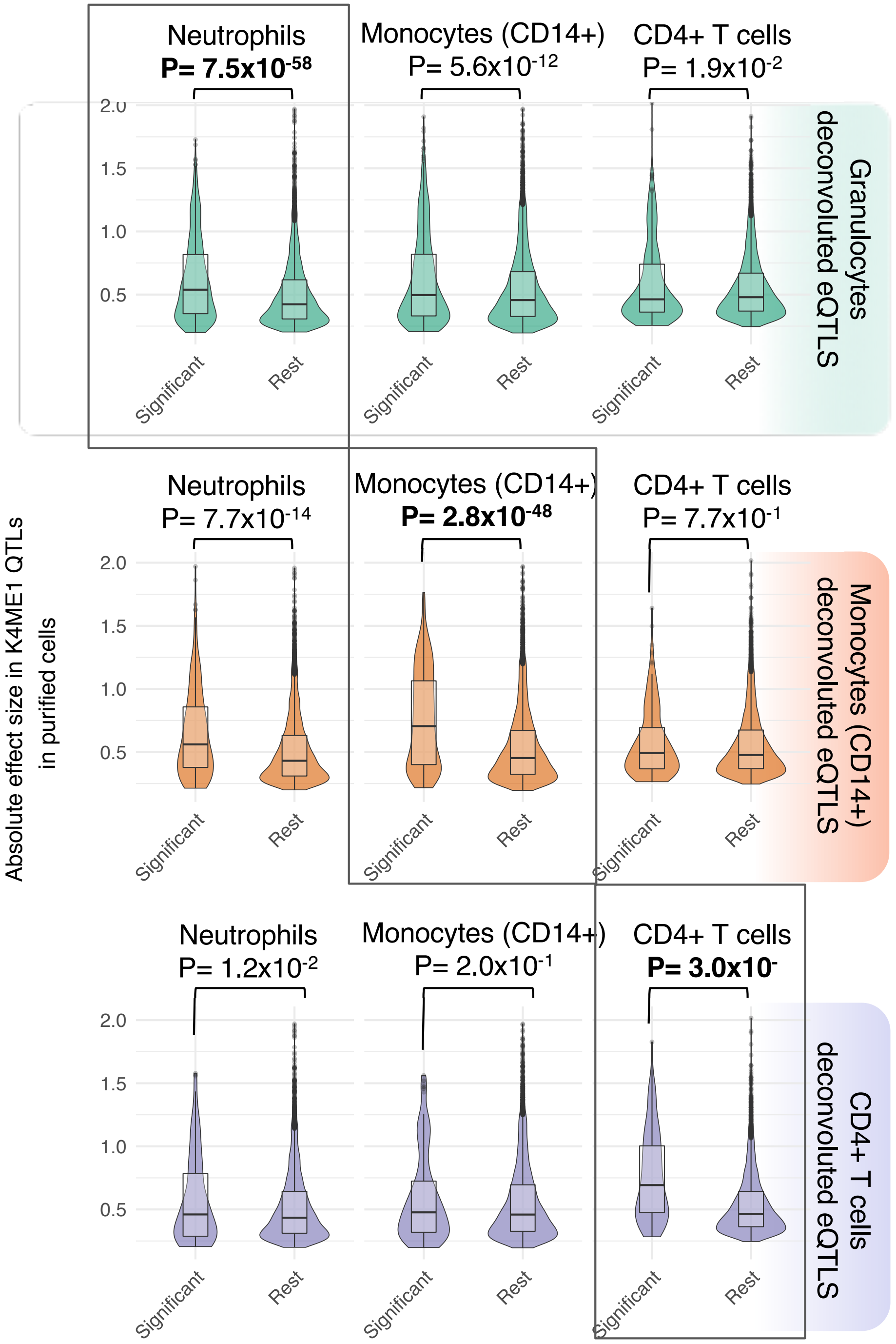


**Supplementary Figure 10. Validation of deconvoluted eQTLs using effect sizes of** K4ME1 **QTLs from purified cells.** Deconvoluted eQTLs (FDR ≤ 0.05) show a significantly bigger effect size for K4ME1 QTLs (where the eGenes is the closest gene tagging the K4ME1 QTLs peak)from their relevant cell subtype compared to the rest of the significant whole blood eQTLs (diagonal boxed comparisons). The off-diagonal comparisons show that these eQTL genes are specific to a cell subtype because the differences in effect sizes are non-significant in all but the comparisons between neutrophils and monocytes (CD14+).

Supp. Figure 10

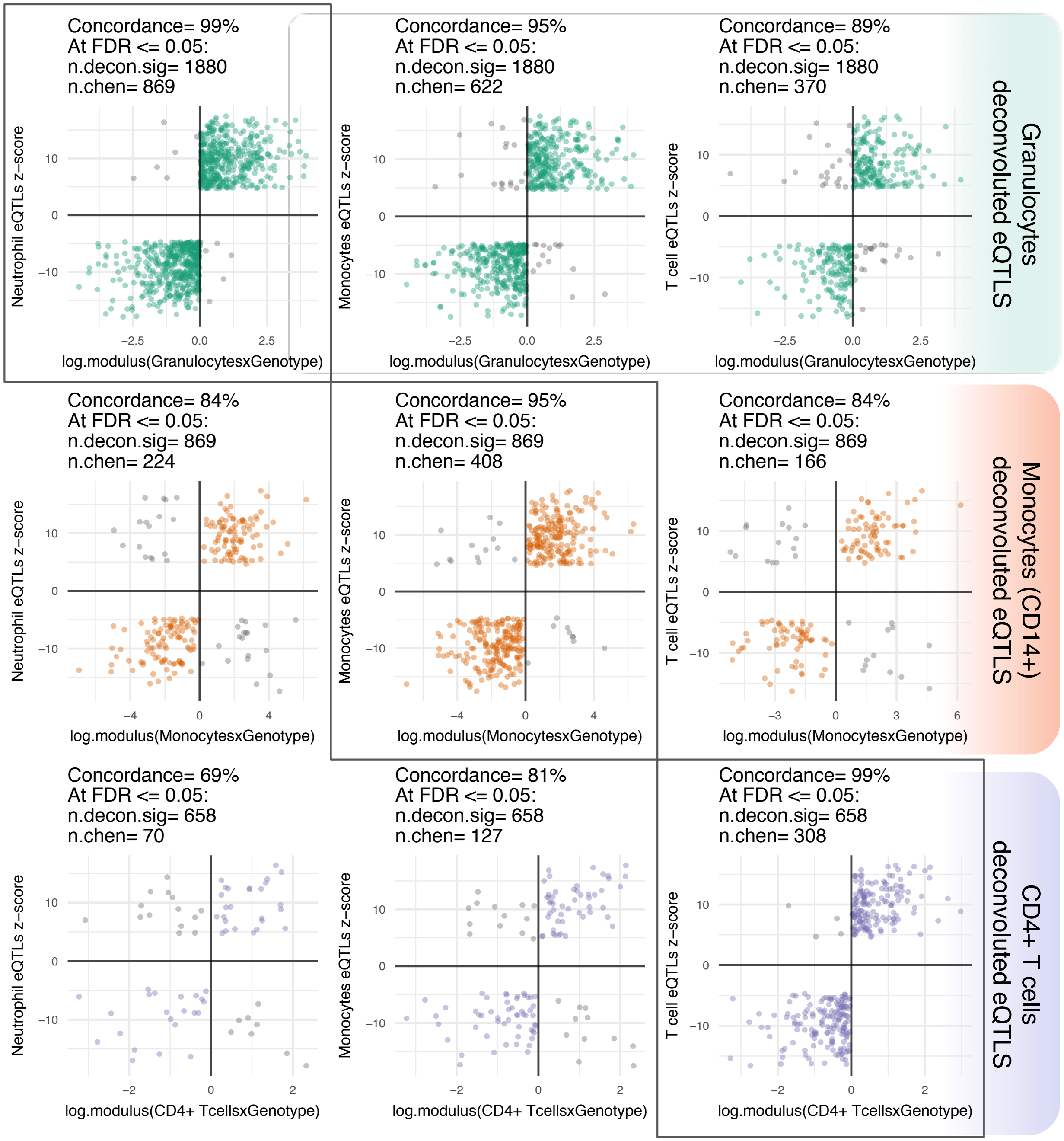


**Supplementary Figure 11. Validation of deconvoluted eQTLs using allelic concordance with eQTLs results from purified cells.** Deconvoluted eQTLs (FDR ≤ 0.05) show a high allelic concordance in their respective purified cell eQTLs. Top row shows allelic concordance of deconvoluted granulocyte eQTLs (all in green) against neutrophils, monocytes and CD4+ T cells. Second row shows deconvoluted monocyte eQTLs against purified cell eQTLs in the same order as top row; bottom row shows the same comparisons as for deconvoluted CD4+ eQTLs. Allelic concordance of the off-diagonal (comparing deconvoluted eQLTs with non-relevant cell types) show a consistent decrease in allelic concordance.

Supp. Figure 11

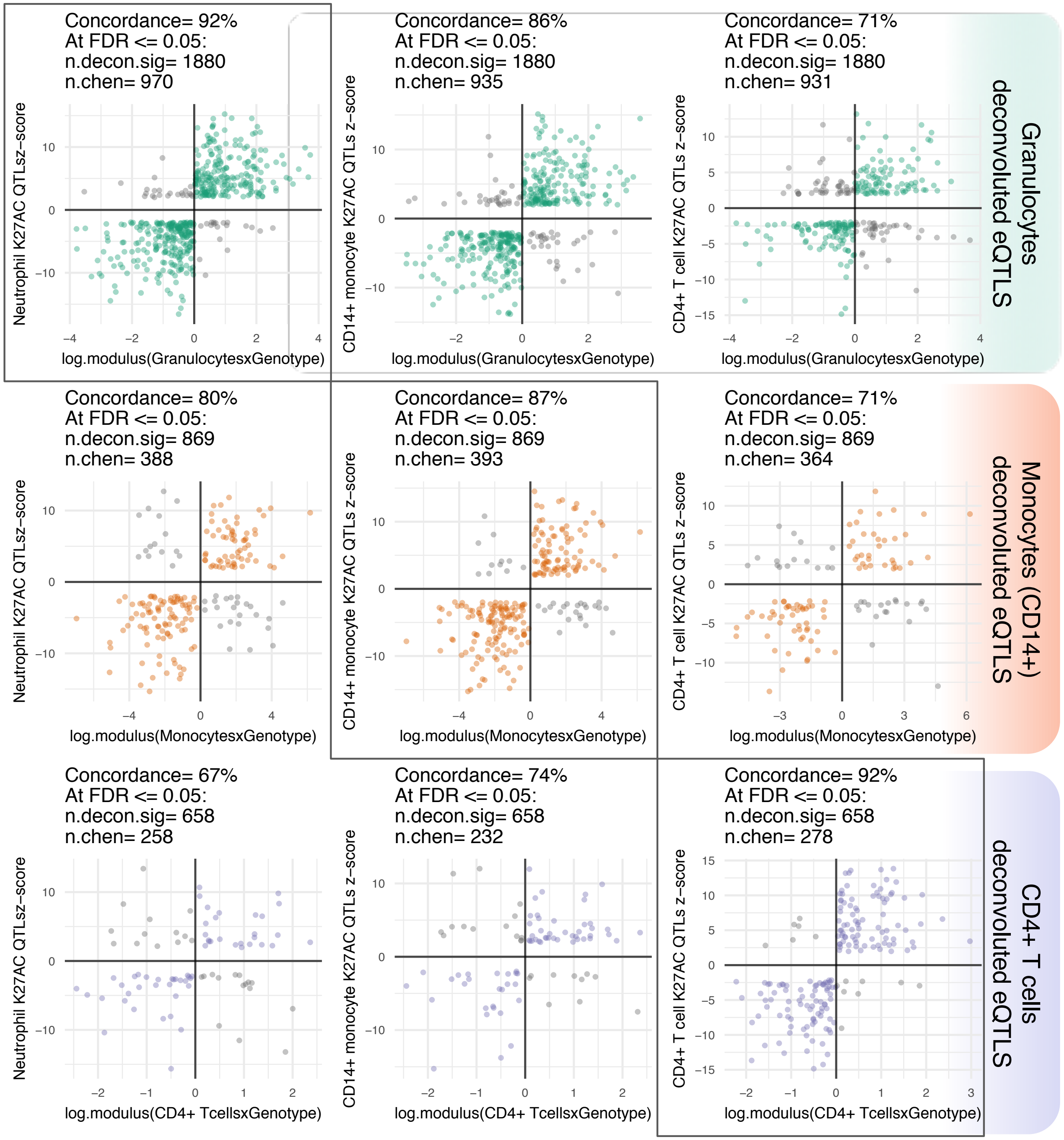


**Supplementary Figure 12. Validation of deconvoluted eQTLs using allelic concordance with K27AC results from purified cells.** Deconvoluted eQTLs (FDR ≤ 0.05) show a high allelic concordance in their respective purified cell K27AC QTLs. Top row shows allelic concordance of deconvoluted granulocyte eQTLs (all in green) against neutrophils, monocytes and CD4+ T cells derived K27AC QTLs. Second row shows deconvoluted monocyte eQTLs (all in orange) against purified cell K27AC QTLs in the same order as top row; bottom row shows the same comparisons as for deconvoluted CD4+ eQTLs (all in purple). Allelic concordance of the off-diagonal (comparing deconvoluted eQLTs with non-relevant cell types) show a consistent decrease in allelic concordance when compared to the relevant cell type comparisons.

Supp. Figure 12

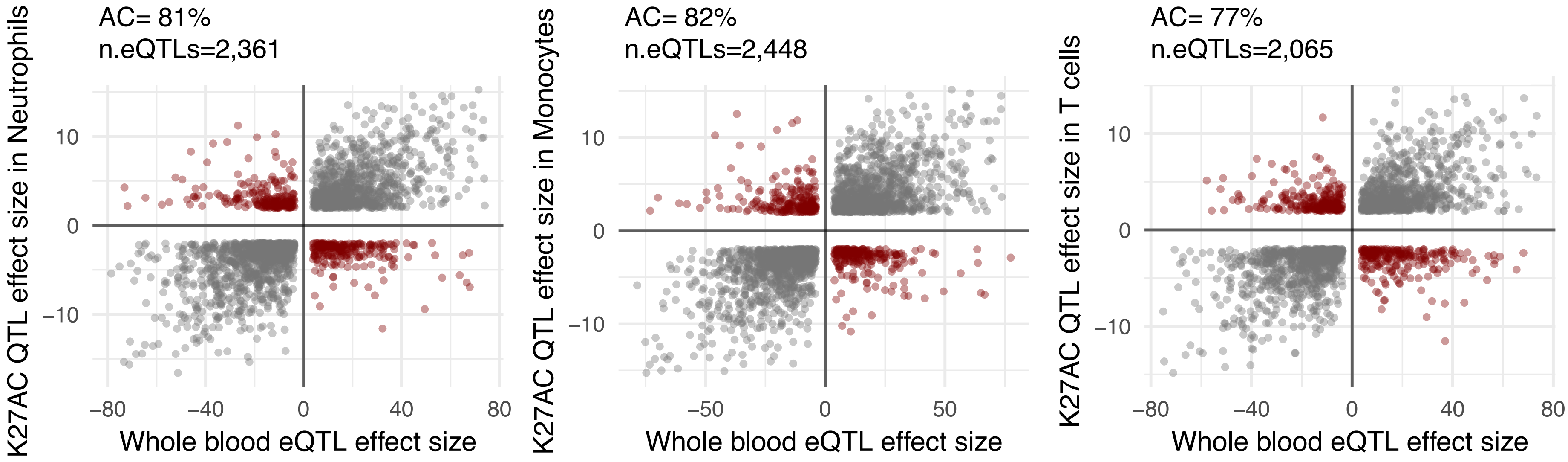


**Supplementary Figure 13. Allelic concordance between whole blood eQTLs and K27AC QTLs** for purified neutrophils, CD14+ monocytes and CD4+ T cells.

Supp. Figure 13

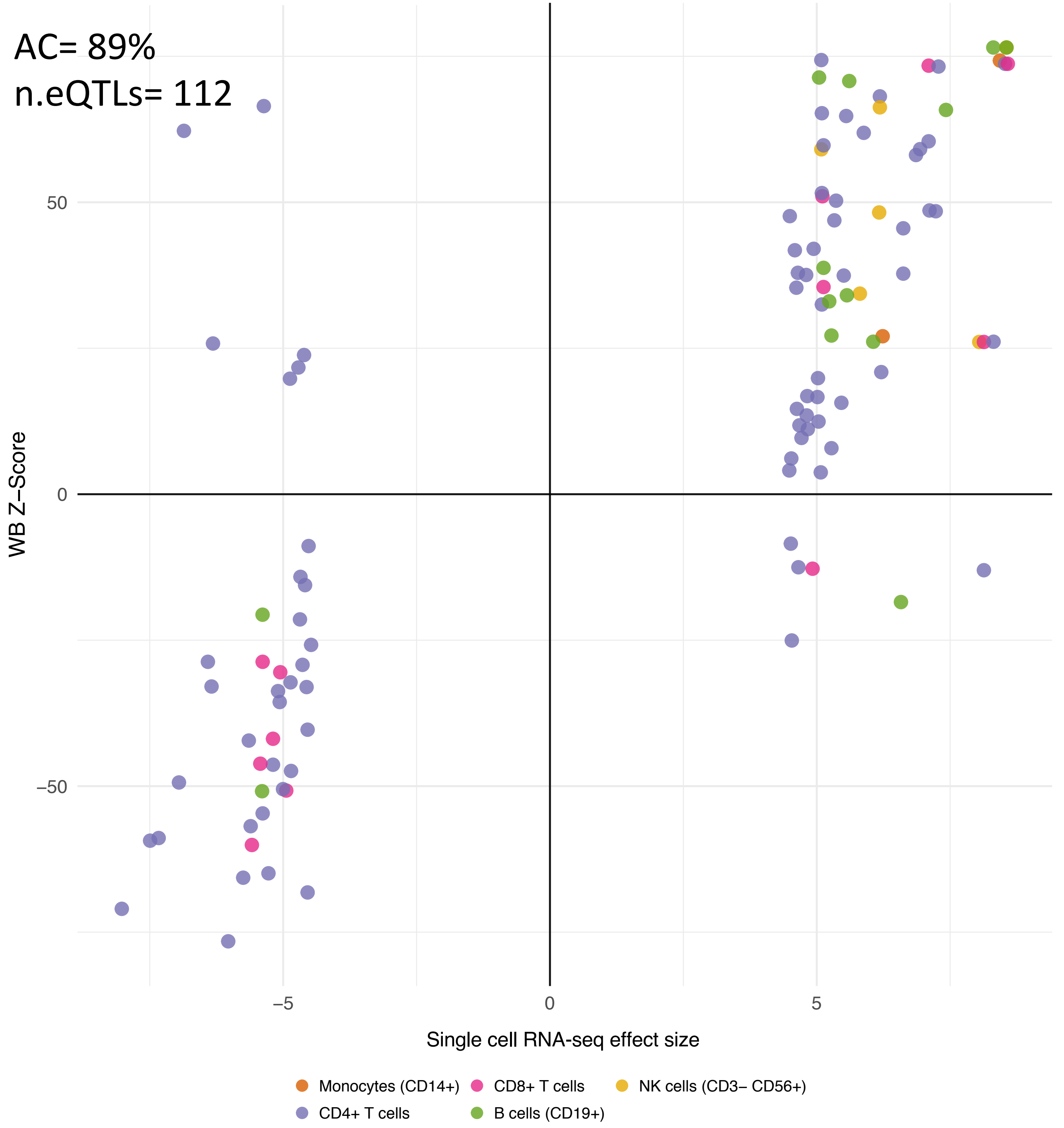


**Supplementary Figure 14. Comparison of whole blood eQTLs with eQTLs from single cell RNA-seq** Whole blood eQTLs show 89% allelic concordance for significant eQTLs derived from single-cell RNA-seq data, comprising monocytes CD14+, B cells, CD4+ T cells, CD8+ T cells and NK cells.

Supp. Figure 14

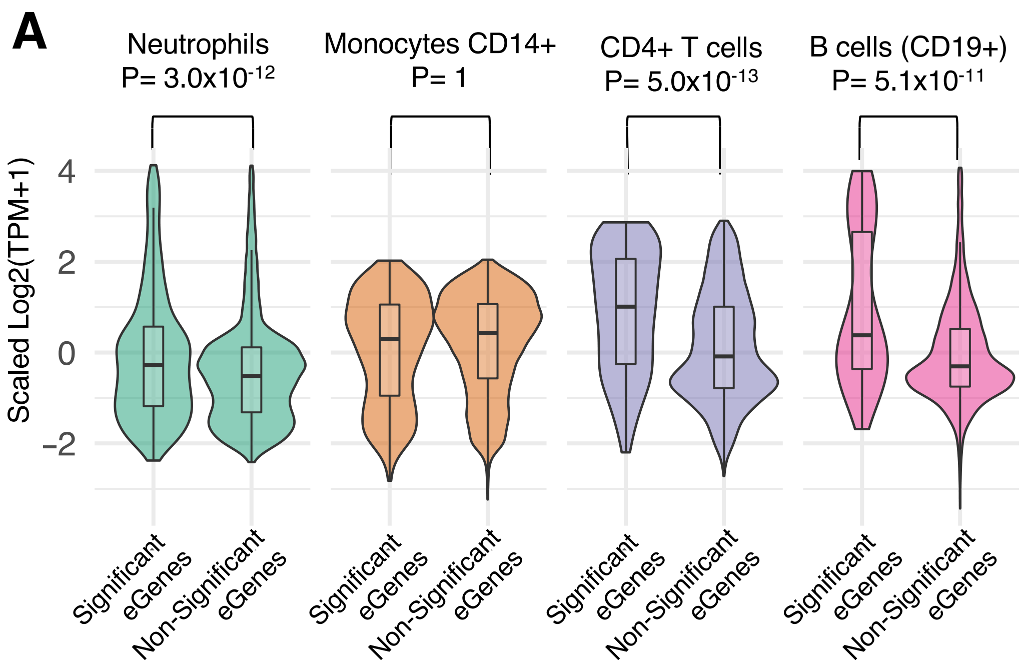


**Supplementary Figure 15. Validation of cell type eQTLs detected in the BIOS cohort using Westra *et at*, method**: (A) Expression of eGenes in purified cell subpopulations from BLUEPRINT (green for granulocyte eQTL genes showing expression for purified neutrophils; orange for monocytes; purple for CD4+ T cells; pink for B cells). (B) CT eQTLs detected by the Westra method show a significantly larger effect size in purified cell eQTLs[11] compared to the rest of the whole blood eQTLs. Boxed-diagonal show the comparisons with relevant cell types, were the effect differences are stronger.

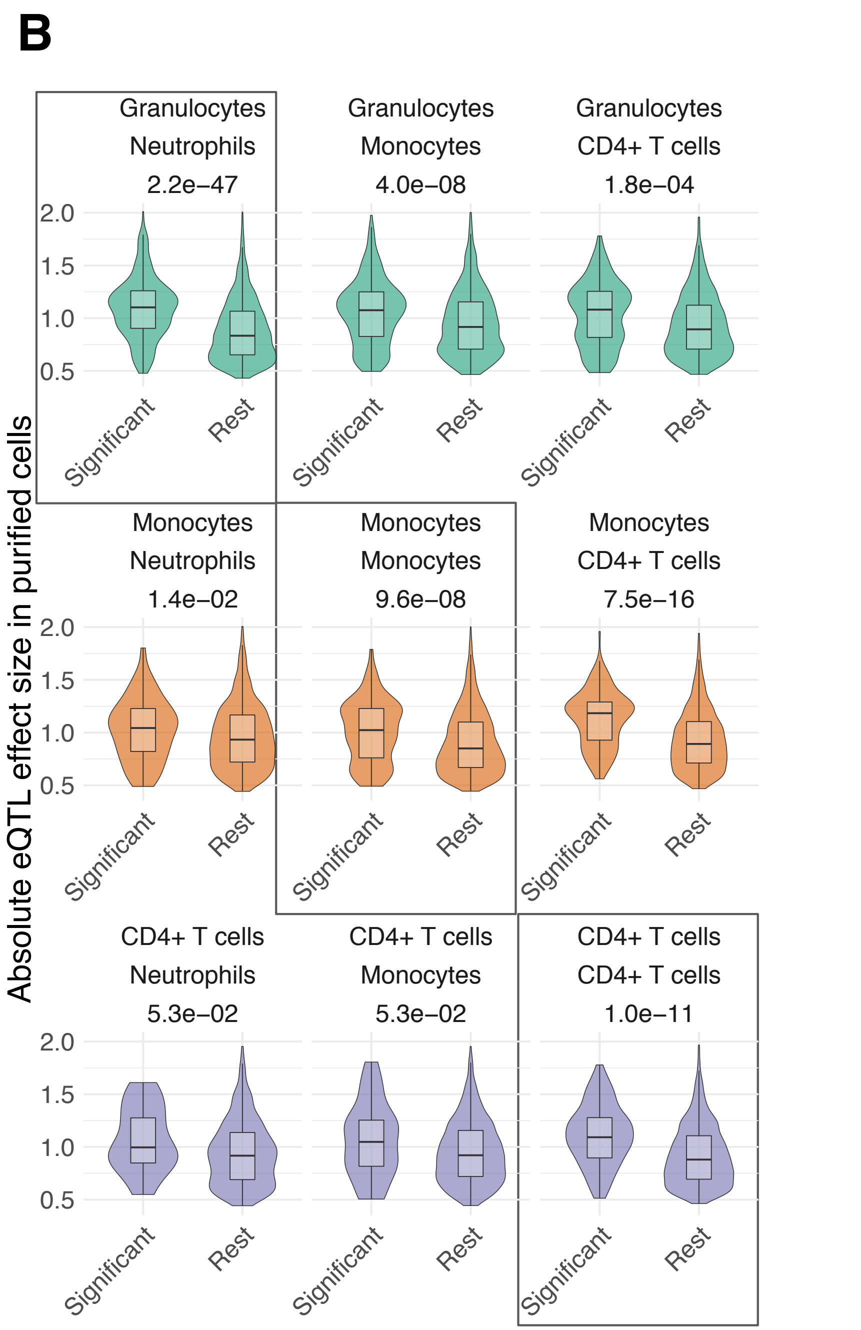

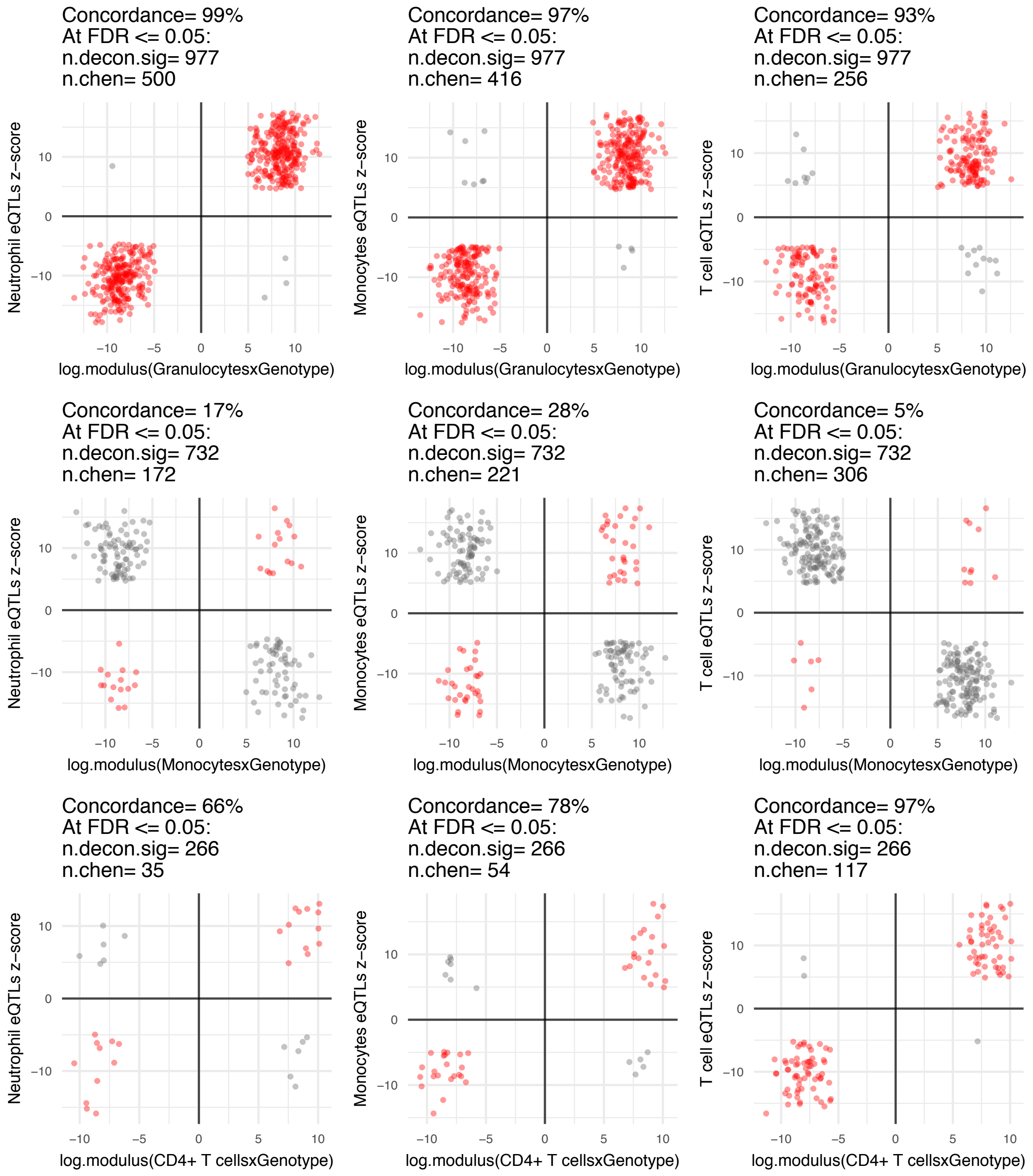


**Supplementary Figure 16. Allelic concordance rates of cell type eQTLs detected using the Westra *et al* method and eQTLs from purified cells**. Top row shows allelic concordance of granulocyte CT eQTLs against neutrophils, monocytes and CD4+ T cells. Second row shows CT monocyte eQTLs against purified cell eQTLs in the same order as top row; bottom row shows the same comparisons for CT CD4+ eQTLs.

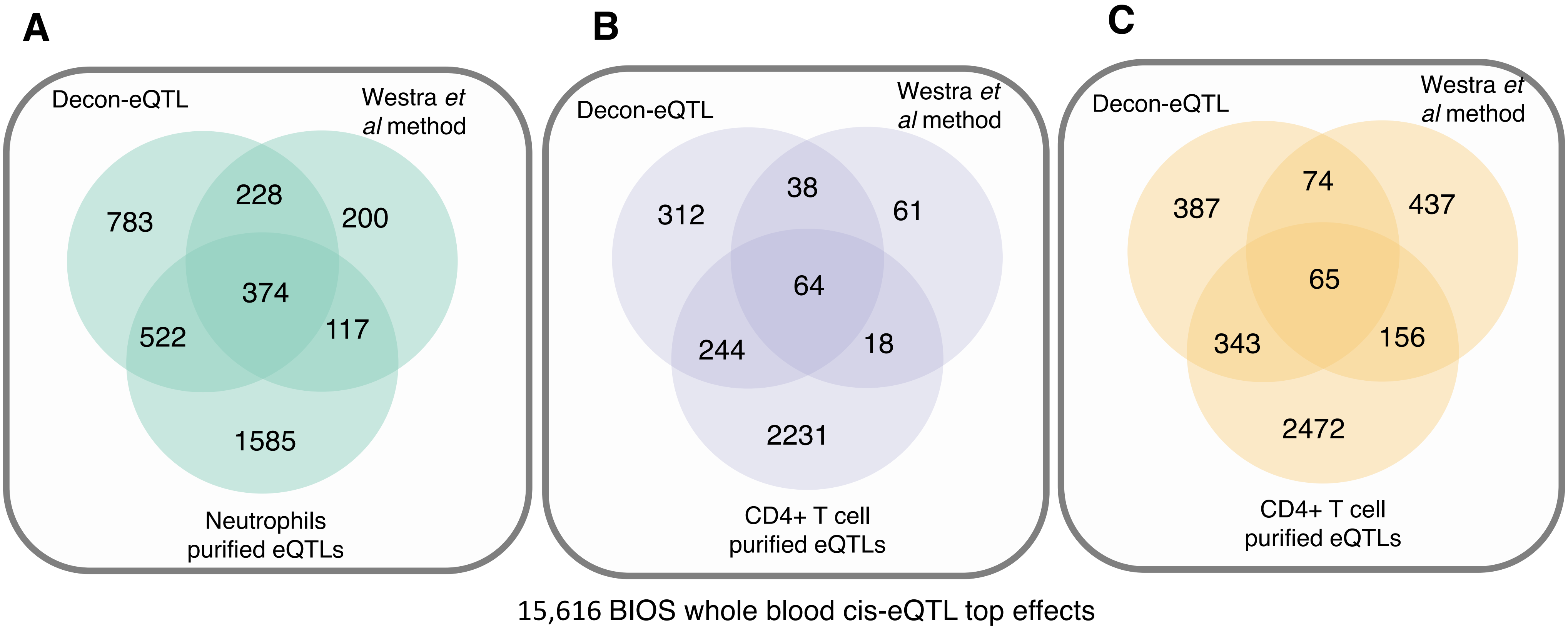


**Supplementary Figure 17. Comparison of Decon-eQTL with Westra *et al* method.** Overlap of CT eQTLS detected with Decon-eQTL, the Westra *et al* method and those found to be significant in purified cell subpopulations, for granulocyte QTLs (A), CD4+ T cells (B), and monocytes (C).

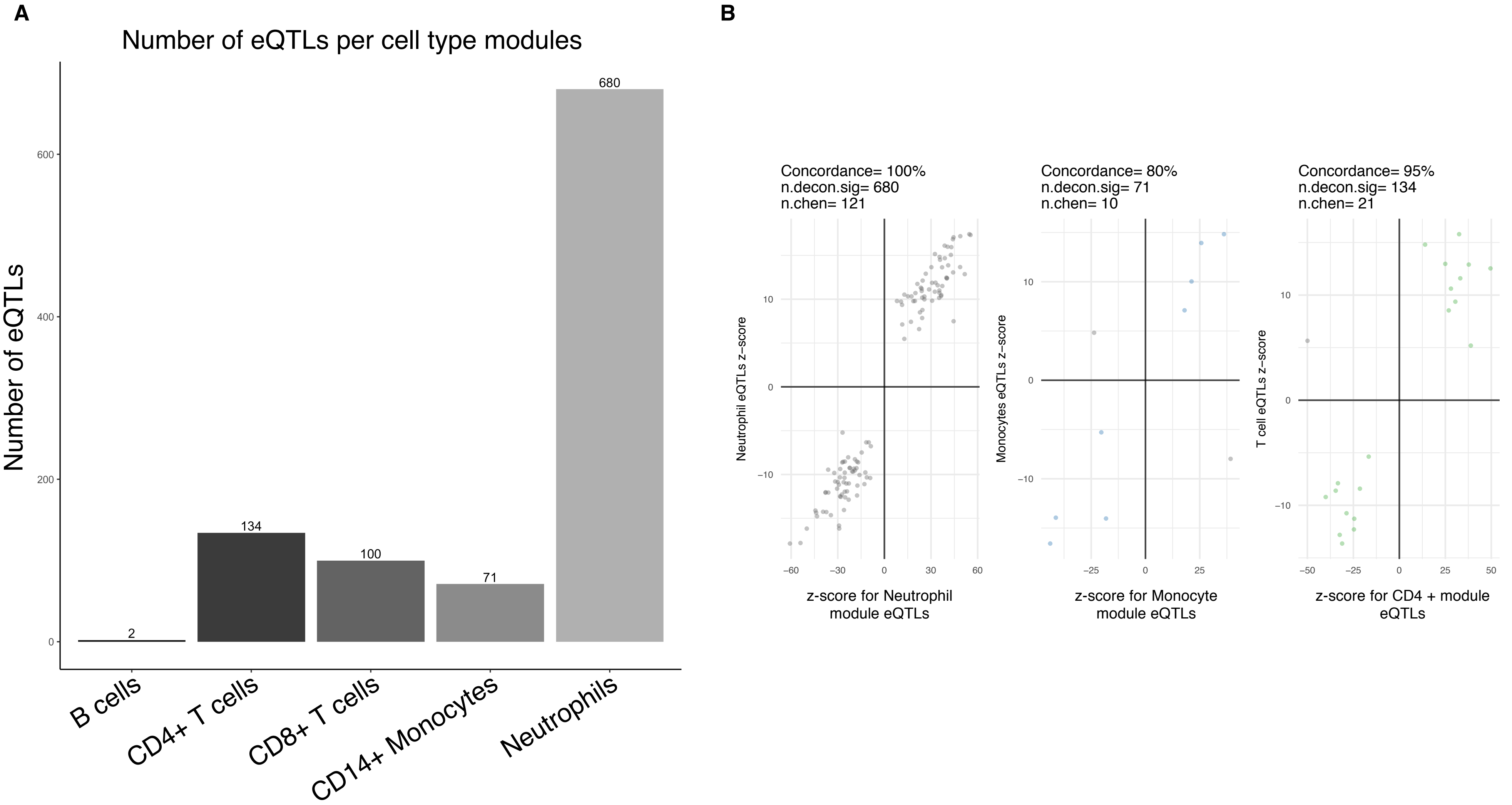


**Supplementary Figure 18. Comparison of Decon-eQTL with other methods for detecting cell type eQTLs**. Total number of eQTLs per cell proportion module obtained by Zhernakova et al. (Nat Gen, 2017) (A). Allelic concordance between overall z-score for eQTLs from neutrophil, monocytes and CD4+ T cell modules against the effect size of purified eQTLs from neutrophils, monocytes and CD4+ T cells.

Supp. Figure 18

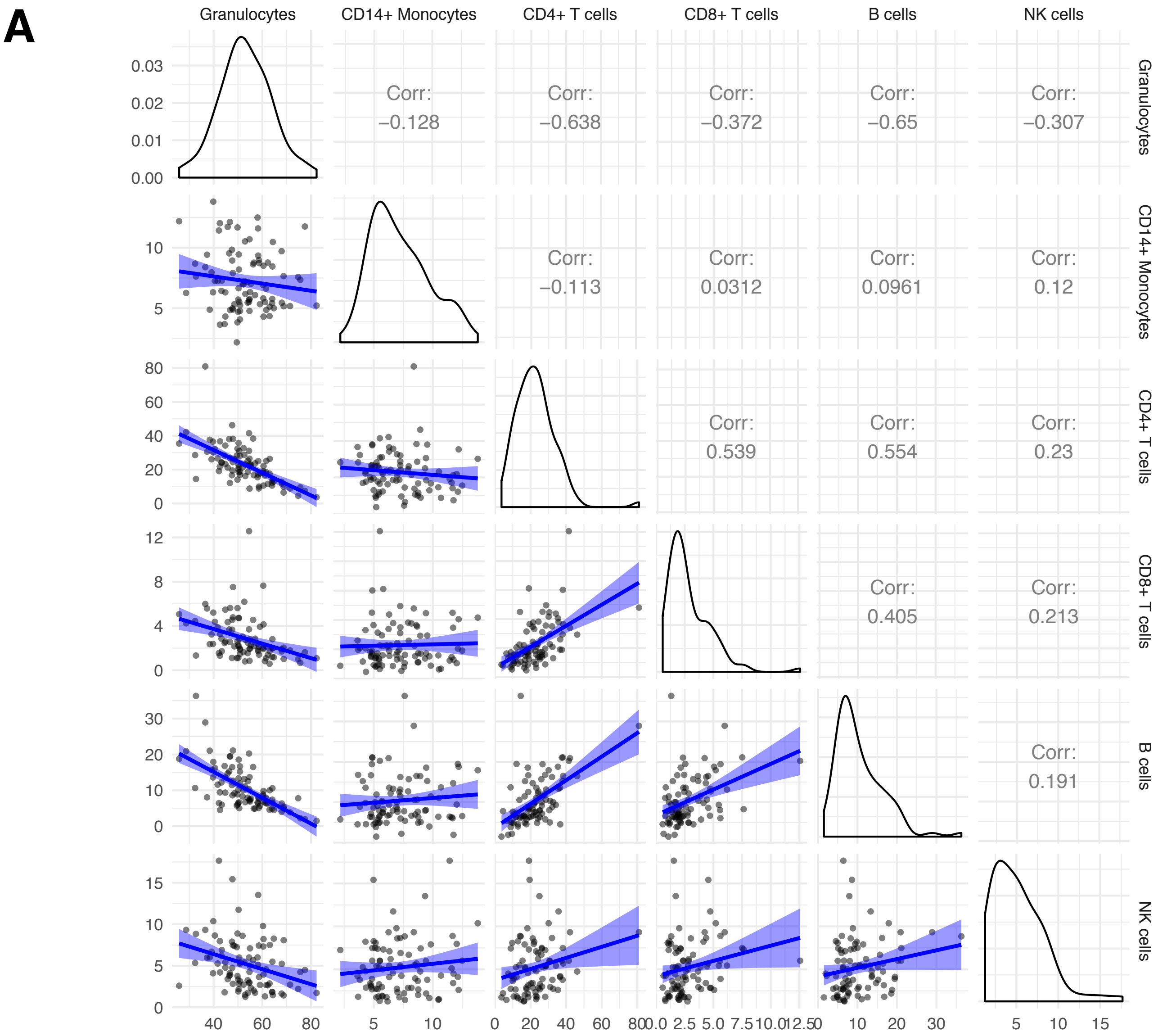


**Supplementary Figure 19. Distribution and correlation among circulating cell proportions.** (A) With 89 samples from 500FG, the scatter plots show the correlations between different cell subpopulations. Blue line indicates a fitted linear model. Diagonal plots depict the overall density distribution per cell type. Upper right triangle shows the Pearson correlation coefficient for each pairwise comparison. (B) shows correlations between different cell subpopulations in the BIOS cohort, which were obtained by prediction using Decon-cell.

Supp. Figure 19A

**B**

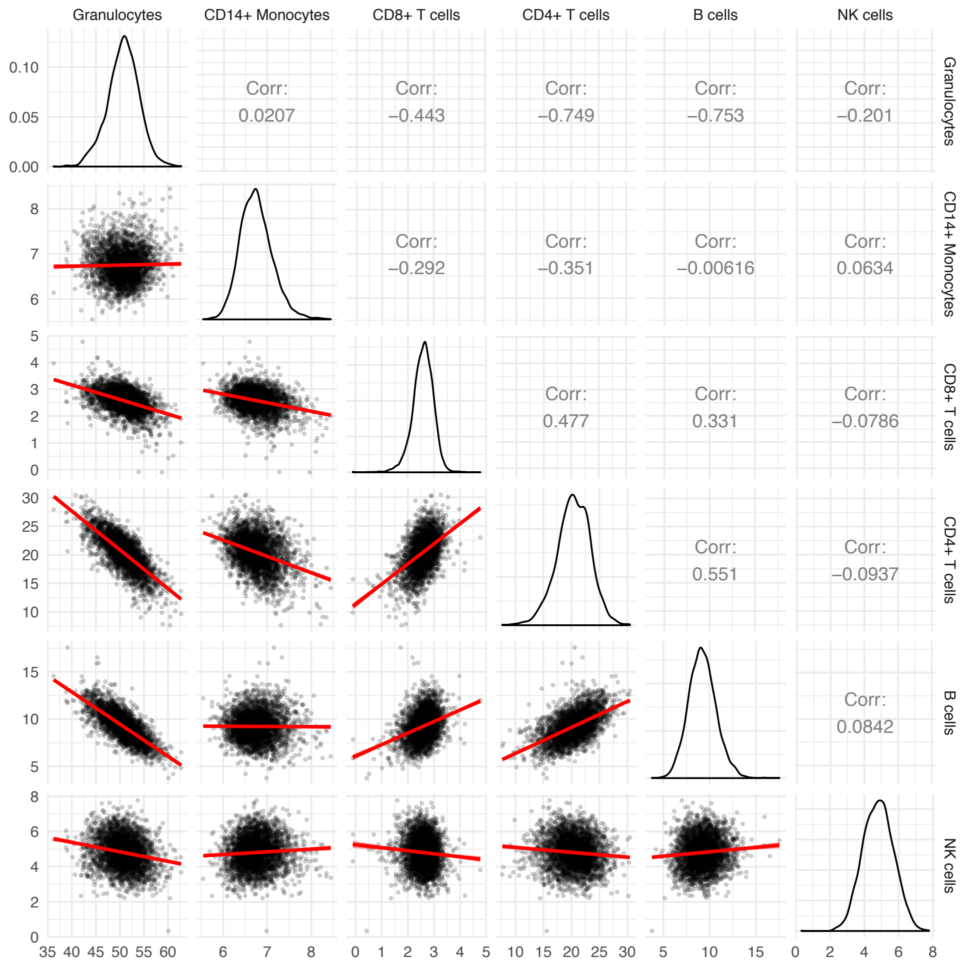


Supp. Figure 19B
